# MOCCS profile analysis clarifies the cell type dependency of transcription factor-binding sequences and cis-regulatory SNPs in humans

**DOI:** 10.1101/2022.04.08.487641

**Authors:** Saeko Tahara, Takaho Tsuchiya, Hirotaka Matsumoto, Haruka Ozaki

## Abstract

Transcription factors (TFs) show heterogeneous DNA-binding specificities in individual cells and whole organisms in natural conditions): *de novo* motif discovery usually provides multiple motifs even from a single ChIP-seq sample. Despite the accumulation of ChIP-seq data and ChIP-seq-derived motifs, the diversity of DNA-binding specificities across different TFs and cell types remains largely unexplored. Here, we propose MOCCS profiles, the new representation of DNA-binding specificity of TFs, which describes a ChIP-seq sample as a profile of TF-binding specificity scores (MOCCS2scores) for every *k*-mer sequence. Using our *k*-mer-based motif discovery method MOCCS2, we systematically computed MOCCS profiles for >10,000 human TF ChIP-seq samples across diverse TFs and cell types. Comparison of MOCCS profiles revealed the global distributions of DNA-binding specificities, and found that one-third of the analyzed TFs showed differences in DNA-binding specificities across cell types. Moreover, we showed that the differences in MOCCS2scores (ΔMOCCS2scores) predicted the effect of variants on TF binding, validated by *in vitro* and *in vivo* assay datasets. We also demonstrate ΔMOCCS2scores can be used to interpret non-coding GWAS-SNPs as TF-affecting SNPs and provide their candidate responsible TFs and cell types. Our study provides the basis for investigating gene expression regulation and non-coding disease-associated variants in humans.

## Introduction

Gene expression regulation is one of the most important mechanisms underlying proper cell function. Dysregulation of gene expression results in diseases such as developmental disorders and cancer. Gene expression is regulated by transcription factors (TFs), which bind to DNA by recognizing sequences specific to each TF. The human genome is estimated to contain more than 1,600 human TFs, and these TFs comprise more than 70 DNA-binding domain types (1). Studies of TF-binding sites have revealed that TFs do not bind to a single particular sequence but rather to a distinct set of similar DNA sequences. Such specific sequence patterns are called DNA-binding motifs. These motifs can be understood using several representations, including consensus sequences, position weight matrices (PWMs), *k*-mers, and hidden Markov models (2–4).

ChIP-seq detects *in vivo* TF-binding sites (TFBSs) in a genome-wide manner and can generate thousands of DNA sequences with lengths of several hundred base pairs from TFBSs. Many specialized tools have been developed for *de novo* motif discovery from ChIP-seq data (5). Usually, more than one single motif is found, even in a single ChIP-seq sample (6). This reflects the heterogeneous sequence context around TFBSs due to ambiguity in DNA recognition, different binding modes (e.g., heterodimerization, cooperative binding, and tethering), and the existence of transcriptional cofactor motifs. Motifs derived by data from ChIP-seq and other assays (e.g., protein binding microarrays and SELEX) have been summarized in several TF motif databases as PWMs or position frequency matrices (7–10).

Despite the accumulation of ChIP-seq data and ChIP-seq-derived motifs, the diversity of TF-binding DNA sequences is largely unknown. In particular, differences in TF-binding sequences among different cell types or different TFs have not been systematically explored. Many systematic studies of large ChIP-seq datasets compared the localization and colocalization of TFBSs among cell types and TFs and revealed cell type specificities in TFBSs (e.g., (11)) and TF regulatory relationships (12). TFBS differences for the same TFs in different cell types have been attributed to changes in the TF partner (13) or epigenome (14). In contrast, several studies attempted to identify discriminative motifs among a small number of ChIP-seq samples and revealed distinct motifs among homologous TFs (15, 16), cooperative partner TFs (17), or different cell types (18). However, the extent of the diversity of TF-binding sequences across different TFs and cell types remains largely unexplored.

Recently, ChIP-seq data have been collected in secondary databases (19–22). These compendiums of ChIP-seq data provide opportunities to analyze the diversity of TF-binding sequences. For this purpose, *k*-mer representation is useful because representations such as PWMs might miss sequences with low frequency (23, 24). Moreover, PWMs cannot evaluate the dependency among base positions. Besides, *k*-mer representation has high interpretability and can easily be transferred to experimental validations such as reporter assays (24, 25). Several methods have been proposed to discover *k*-mer motifs in ChIP-seq data (4, 23, 24, 26–30) or to predict the effect of nucleotide substitutions on TF binding (31). Thus, comprehensive analysis and comparison of *k*-mer representations of the TF-binding sequences of each ChIP-seq sample would reveal the diversity of TF-binding sequences among different cell types and TFs. This approach is exemplified by studies comparing *k*-mer motifs among different TFBSs of the same TF (29) or among homologous TFs (16).

Here, we propose MOCCS profiles, the new representation of DNA-binding specificity of TFs, which describes a ChIP-seq sample as a profile of TF-binding specificity scores (MOCCS2scores) for every k-mer sequence. Using our *k*-mer-based motif discovery method MOCCS2, we systematically computed MOCCS profiles for >10,000 human TF ChIP-seq samples across diverse TFs and biological contexts (tissue and cell types). By comparing the MOCCS profiles across high-quality ChIP-seq samples, we confirmed that similarities in TF-binding sequences were marked by similarities in cell types (cell type classes) and TFs (TF families) and interactions with other TFs. We also found that a third of the analyzed TFs showed differences in DNA-binding specificities across cell types. Moreover, differential analysis of the MOCCS profiles revealed differentially bound *k*-mers between different cell types and different TFs. Furthermore, we showed that differences in MOCCS2scores (ΔMOCCS2scores) can predict the effects of variants on TF binding, which were validated with the results of *in vitro* and *in vivo* assays. We also showed that the ΔMOCCS2score can be used to interpret non-coding GWAS-SNPs as TF-affecting singlenucleotide variants and associate them with candidate responsible TFs and cell types. Our study demonstrates the MOCCS profile analysis provides the basis for investigating gene expression regulation and non-coding disease-associated variants in humans.

## Results

### Overview of the MOCCS profile

To obtain a collection of TF-binding sequences as *k*-mers and elucidate the diversity of TF-binding sequences, we applied MOCCS2 (23, 29) to the 10,534 peak calling results (human, hg38) from the ChIP-seq data repository ChIP-Atlas (21) (**Fig. 1A**). MOCCS2 quantifies the binding specificity of each *k*-mer as the MOCCS2score of a ChIP-seq sample. We refer to the vector of MOCCS2scores for all *k*-mers as the MOCCS profile of the ChIP-seq sample. A high MOCCS2score indicates high binding specificity. For example, when MOCCS2 was applied to the GATA3 ChIP-seq sample from the MDA-MB-231 cell line (breast), the *k*-mer AGATAA, which was supported by the known GATA3 PWM motif, showed the highest MOCCS2score (**Fig. 1B**).

**Figure 1.**
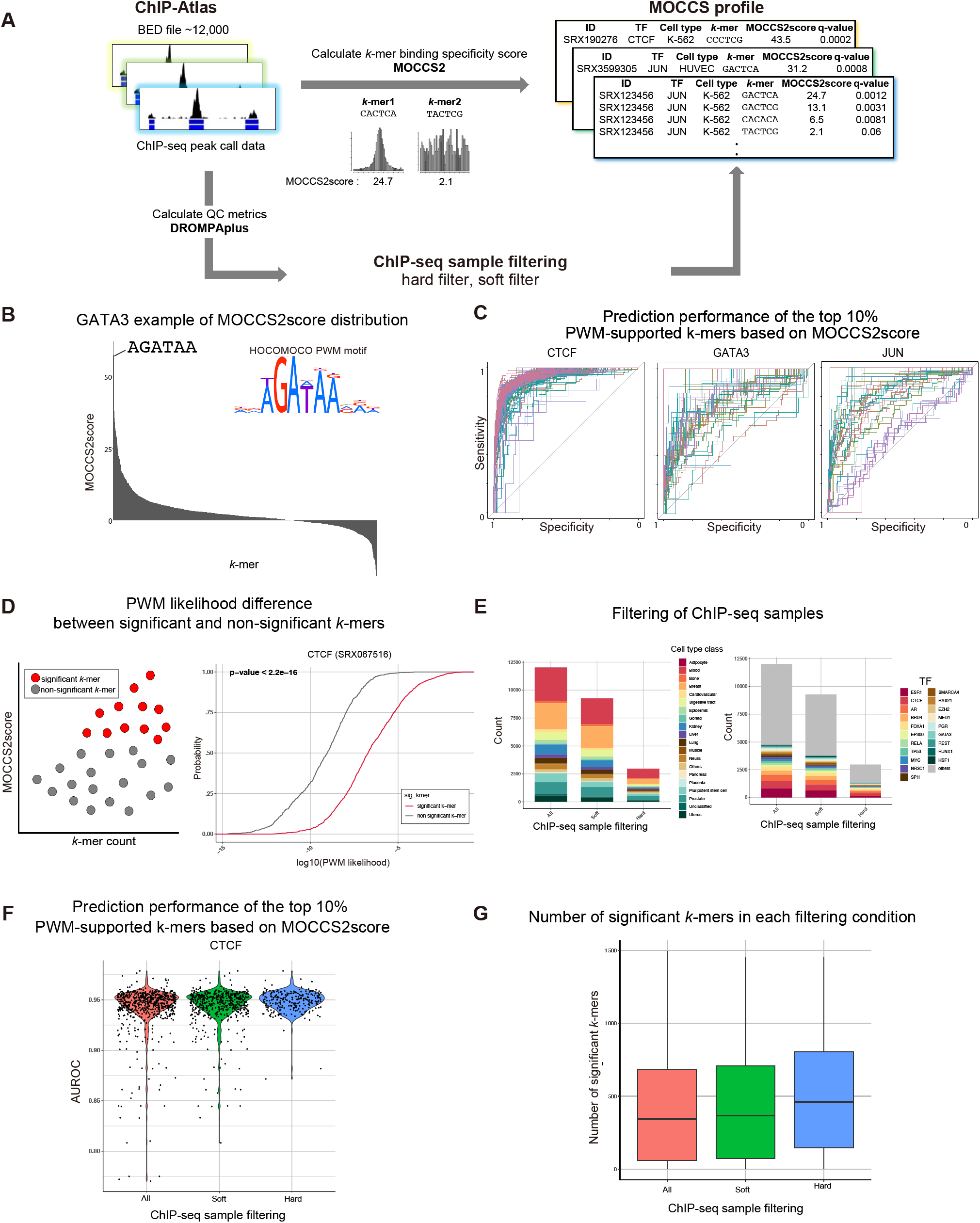
Overview of the MOCCS profile. **A**: Schematic overview of the ChIP-seq data processing. MOCCS2 was applied to BED files of human ChIP-seq samples from ChIP-Atlas, resulting in MOCCS profiles, containing *k*-mers, MOCCS2scores, q-values, and annotations. In parallel, quality control metrics for ChIP-seq samples were calculated for filtering samples (hard and soft filters). **B**: An example of a MOCCS profile for a GATA3 ChIP-seq sample of MDA-MB231. The *k*-mer with the highest MOCCS2score (AGATAA) was similar to the GATA3 PWM motif from the HOCOMOCO database. **C**: Prediction specificity and sensitivity for the top 10% PWM-supported *k*-mers based on the MOCCS2score. Each line represents a different ChIP-seq sample. **D**: Significant *k*-mers are supported by PWMs. Left: Schematic overview of significant *k*-mer detection, and the significant *k*-mers defined by q-values contain PWM-supported *k*-mers. Right: Cumulative relative frequencies of the PWM likelihood for the significant *k*-mers (q < 0.05) and the other *k*-mers of a CTCF ChIP-seq sample. **E**: The bar plots show the number of ChIP-seq samples passing soft- and hard-filtering. Bars are colored by cell type class (left) or TF (right). **F**: Violin plot showing the AUROC of the prediction of the top 10% PWM-supported *k*-mers based on the MOCCS2score. The red violin plot is all of the CTCF ChIP-seq samples, the green plot is the soft-filtered CTCF ChIP-seq samples, and the blue plot is the hard-filtered CTCF ChIP-seq samples. High-quality ChIP-seq samples with high AUROC scores remaining after soft and hard filters. **G**: Distribution of the significant *k*-mers in ChIP-seq samples that passed each filtering step.

To further confirm that a high MOCCS2score is indicative of TF-binding sequences, we associated the MOCCS2score with the likelihood of the known PWMs for *k*-mers. If a *k*-mer with a high MOCCS2score is a TF-binding sequence, the likelihood of the known PWM for the *k*-mer should also be high. We defined a *k*-mer with a top 10% PWM likelihood to be a PWM-supported *k*-mer and evaluated the ability of the MOCCS2score to discriminate PWM-supported *k*-mers from other *k*-mers. Most CTCF (95%), GATA3 (39%), and JUN (52%) ChIP-seq samples showed an area under the receiver operating characteristic curve (AUROC) exceeding 0.8 (**Fig. 1C**). This indicated that the MOCCS profile could detect TF-binding sequences.

To evaluate the statistical significance of MOCCS2scores, we devised a method for calculating the p-value of the MOCCS2score for each *k*-mer and subsequently calculated the q-value for multiple testing correction (**Materials and Methods**). We defined a *k*-mer satisfying a q-value < 0.05 as being a significant *k*-mer. We verified that the q-values of the MOCCS2score showed high performance in the detection of TF-binding sequences (sensitivity > 86.6%, specificity > 99.7%) and effectively controlled for the false-discovery rate (FDR) by using simulated datasets (**Fig. S2, Materials and Methods**). To test whether this q-value could select the *k*-mers recognized by TFs using real ChIP-seq data, we divided the *k*-mers into significant and non-significant *k*-mers and calculated the likelihood of PWM motifs in each set. The PWM likelihood was significantly higher with significant *k*-mers than with non-significant *k*-mers (p < 2.2e-16, Kolmogorov–Smirnov test) (**Fig. 1D**). These results indicated that this q-value can select the *k*-mers significantly recognized by TFs.

We obtained 10,534 MOCCS profiles comprising ChIP-seq samples of 940 TFs and 20 cell type classes. To select high-quality ChIP-seq samples, we first obtained the quality metrics by applying DROMPAplus, a ChIP-seq data quality control tool (32), to the ChIP-seq samples and by downloading processing logs in ChIP-Atlas. Based on the quality metrics, we prepared two types of filters, a soft filter and a hard filter (**Materials and Methods**), and 88.1% and 28.2% of the samples passed each of these filters, respectively (**Fig. 1E, Fig. S1**). To evaluate the improvement with the two filters, we classified *k*-mers with the top 10% PWM likelihoods by the MOCCS2score. The soft and hard filters removed ChIP-seq samples with a low AUROC, indicating that the two filters selected high-quality ChIP-seq samples (**Fig. 1F**). The average number of significant *k*-mers was 439.7 (23.7% of all *k*-mers in a ChIP-seq sample) with the soft filter and 511.5 (27.6%) with the hard filter (**Fig. 1G, Fig. S3**). Collectively, our MOCCS profile dataset comprised vectors of significantly recognized TF-binding *k*-mers with q-values.

MOCCS profile comparisons reveal TF- and TF family-dependent similarities of binding sequences Using the high-quality MOCCS profiles that passed the hard filter, we next compared the MOCCS profiles from different ChIP-seq samples (**Fig. 2A**). In these analyses, which focused on the TF family-dependent similarity of binding sequences (6), we added the annotation of the TF family from the CIS-BP database to the sample metadata (8).

**Figure 2.**
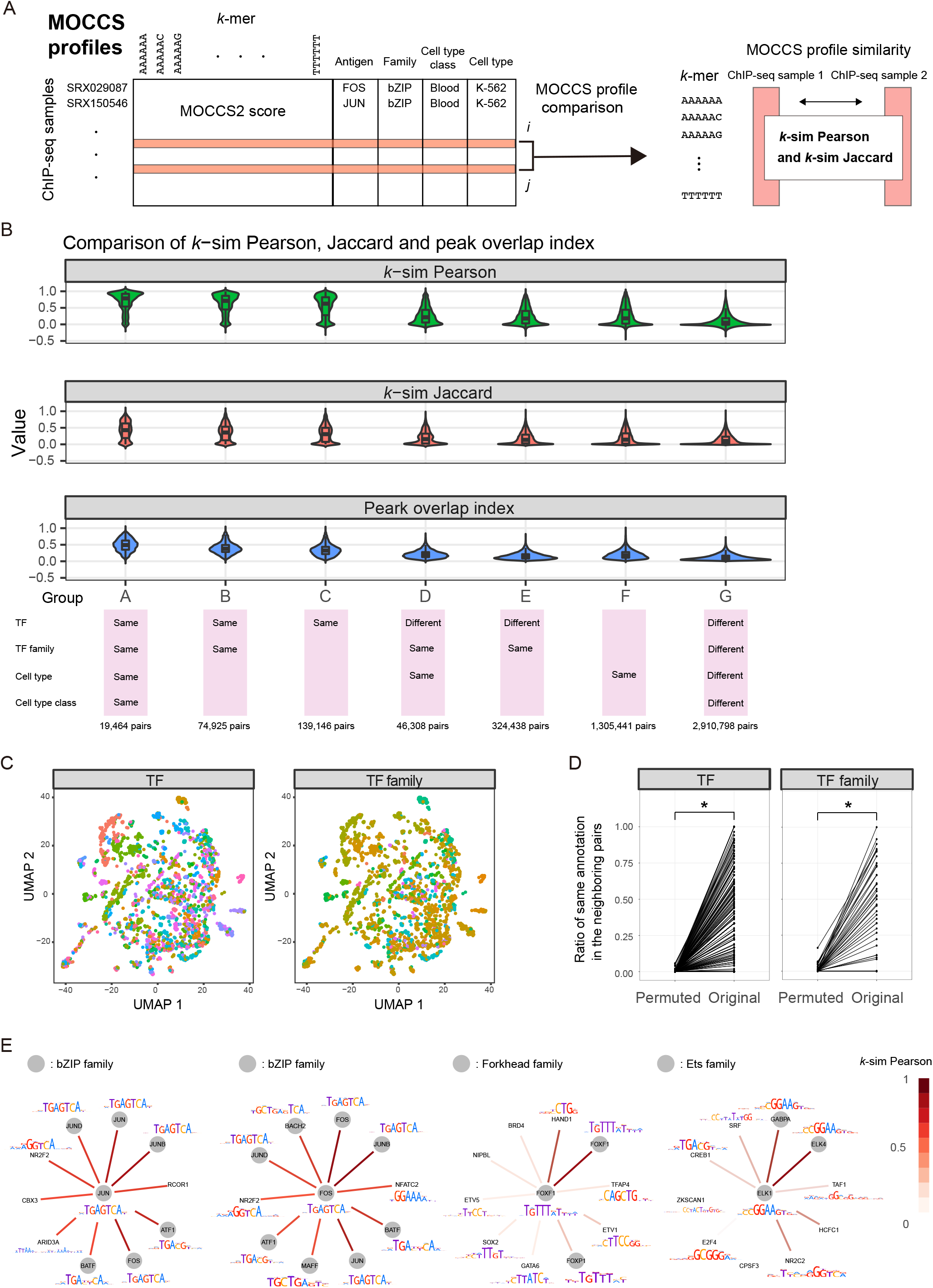
MOCCS profile comparisons reveal the TF- and TF familydependent similarities of binding sequences. **A**: Schematic overview of MOCCS profile comparisons between two ChIP-seq samples. **B**: Comparison of the *k*-mer similarity indices (*k*-sim) Pearson and Jaccard and the peak overlap index. The y-axis indicates the value of each index. Each group is arranged along the x-axis. The color indicates the type of index. **C**: UMAP visualization of the MOCCS profiles. Each dot indicates each ChIP-seq sample. The color indicates the TFs or TF families. **D**: Ratios of neighboring pairs with the same annotations (left, TF; right, TF family) for the original and permuted data. Asterisks indicate the statistical significance between the original and permuted data (non-parametric test, p < 6.26e-249) **E**: TF similarity patterns by the weighted bipartite graph. Each node represents a TF, and the node in the center represents the query TF. For each query TF, the top ten TFs were selected for the *k*-sim Pearson. The color of the edge indicates the *k*-sim Pearson. The color around the nodes indicates the TF family of the query TFs. The obtained PWM motifs from the HOCOMOCO database were indicated. Note that the PWM motifs were not shown in case of not found in the HOCOMOCO database.

To quantitatively compare two MOCCS profiles, we calculated two similarity indices for MOCCS profiles: Pearson (*k*-sim Pearson) and Jaccard (*k*-sim Jaccard) (**Fig. 2A, Materials and Methods**). In addition, we calculated the peak overlap index by using the BED files as a control, which directly reflects the degree of the peak overlap regions (**Materials and Methods**).

To validate our *k*-sim Pearson and Jaccard, we compared the *k*-sim Pearson and Jaccard and the peak overlap index. Based on the annotation match for TF, TF family, cell type class, or cell type, we divided the MOCCS profiles into groups with various combinations of annotation matches (**Fig. 2B**). The *k*-sim Pearson and Jaccard and the peak overlap index consistently showed higher median values in the order of F to A (**Fig. 2B**). Group A, whose annotations were all the same, showed the highest median values, which was consistent with the expectation that the same TFs of the same cell type would have similar MOCCS profiles (**Fig. 2B**). Group G, whose annotations were all different, exhibited the lowest median values, which was consistent for the opposite reason (**Fig. 2B**).

Compared with group G, groups A–F showed a significant increase in the *k*-sim Pearson and Jaccard and in the peak overlap index (Mann–Whitney U test, p < 2.2e-16) (**Fig. 2B**). The comparisons of groups C, E, and F with G revealed significant increases in the *k*-sim Pearson and Jaccard and in the peak overlap index with the same TF, TF family, or cell type class (Mann–Whitney U test, p < 2.2e-16) (**Fig. 2B**). The comparisons of groups B and C and of groups D and E indicated significant increases with the same cell type class, even for the same TF or TF family (Mann–Whitney U test, p < 2.2e-16). Taken together, the *k*-sim Pearson and Jaccard suggested TF-, TF family-, and cell type-dependent similarities in the MOCCS profiles. We illustrate the other comparisons in **Figure S4A**.

We then compared the *k*-sim Pearson and Jaccard with the peak overlap index in each group, revealing significant correlation of the *k*-sim Pearson and Jaccard with the peak overlap index (**Fig. S4B and C**). Accordingly, our *k*-sim Pearson and Jaccard approaches quantify the similarities in TF-binding sequences between two MOCCS profiles in a manner consistent with the peak overlap index.

Next, based on significant increases in the *k*-sim Pearson and Jaccard with the same TF, we assumed the existence of a TF-dependent similarity. We applied UMAP visualization to the MOCCS profiles using *k*-sim Pearson and found adjacency of ChIP-seq samples with the same TF or TF family, supporting the existence of TF-dependent similarities in binding sequences (**Fig. 2C**). This tendency was diminished when we permuted the TF or TF family annotations of ChIP-seq samples, indicating that ChIP-seq samples with the same TF or TF family had similar MOCCS profiles (**Fig. 2D**). Using the *k*-sim Pearson results and by dividing them into the top three pairs and the other pairs, we also confirmed the statistical significance in a chi-square test (**Fig. S5A and B**). Thus, the MOCCS profile comparisons revealed TF-dependent similarities in binding sequences.

Based on the significant increase in the *k*-sim Pearson and Jaccard for pairs of ChIP-seq data with the same TF family, we also hypothesized that the *k*-sim Pearson could extract TF similarity patterns among TFs. From the families with the top ten adjacency values, we selected JUN, FOS, FOXF1, and ELK1 for the demonstrations. We extracted the top 10 similar TFs based on the *k*-sim Pearson for each TF queried (**Fig. 2E**). JUN and FOS extracted the same bZIP family TFs in 14 of the 20 TFs. FOXF1 and ELK1 extracted most TFs with the same TF family in the top three TFs. Furthermore, FOS, JUN, and JUNB were commonly extracted as the top three TFs in JUN and FOS, whose cooperative roles as AP-1 proteins are well-known (33), demonstrating the ability of the *k*-sim Pearson to extract the TF similarity patterns. Collectively, the MOCCS profile comparisons revealed the TF similarity patterns among the TFs and the TF family-dependent similarities in the binding sequences.

### MOCCS profile comparison reveals cell type-dependent TFs and TF similarity patterns

We further investigated the cell type-dependent similarities in the MOCCS profiles (**Fig. 3A**). Again, we performed UMAP visualization and annotated the cell type classes using color, revealing the adjacency of ChIP-seq samples of the same cell type class (**Fig. 3B**). Similar to TF or TF family, this tendency was diminished when we permuted the cell type class annotations of ChIP-seq samples, indicating that ChIP-seq samples with the same cell type class had similar MOCCS profiles (**Fig. 3C**). The statistical significance was also confirmed by chi-square testing between the top three pairs and the other pairs (**Fig. S5C**). Accordingly, MOCCS profile comparisons revealed the existence of cell type-dependent similarities in the binding sequences.

**Figure 3.**
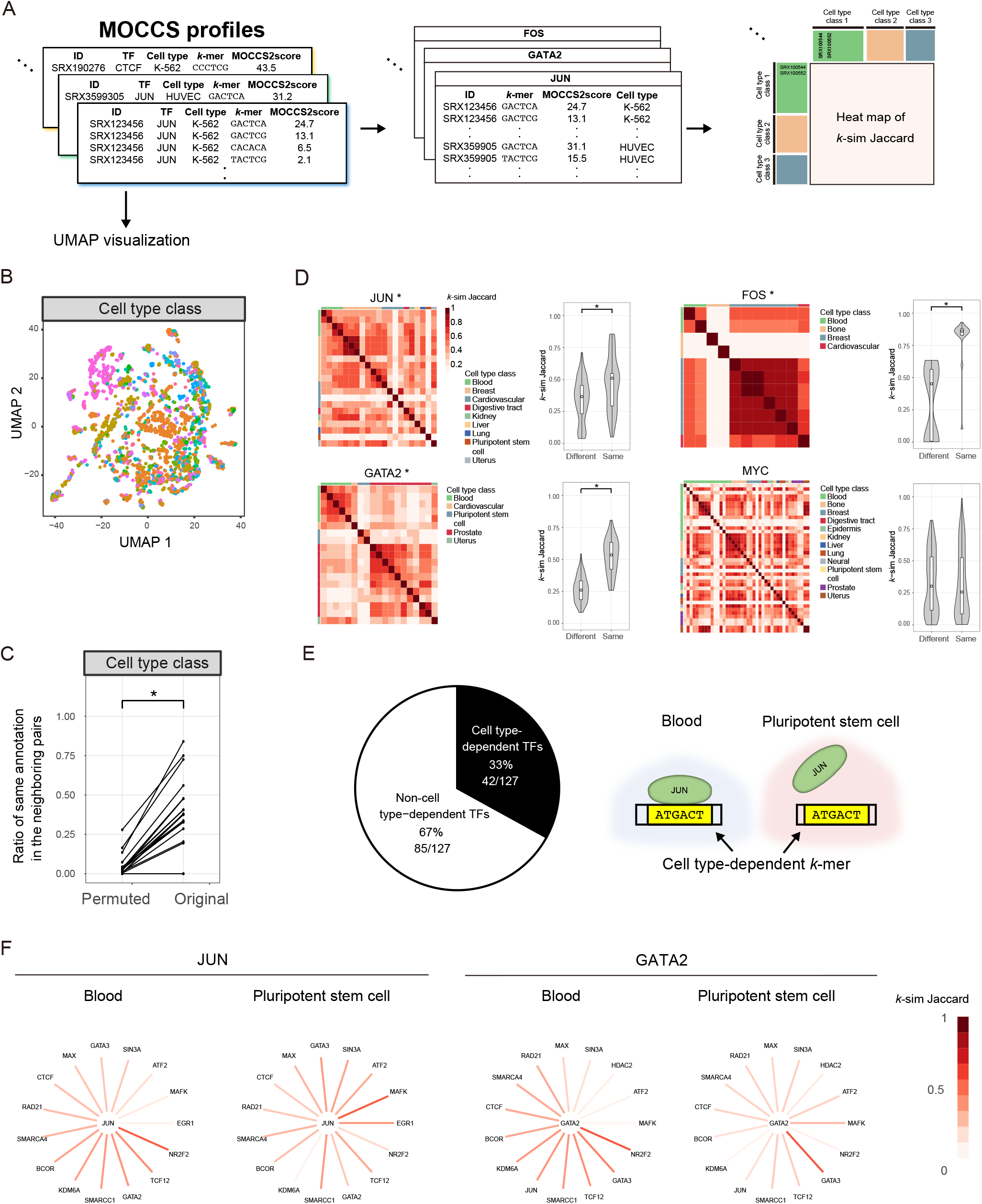
MOCCS profile comparisons reveal cell type-dependent TFs and TF similarity patterns. **A**: Schematic overview of the MOCCS profile comparisons between ChIP-seq samples with the same TF from different cell type classes. **B**: UMAP visualization of the MOCCS profiles. Each point indicates each ChIP-seq sample. The colors of the points represent the cell type classes. **C**: Ratios of neighboring pairs with the same annotations (cell type class) for the original and permuted data. Asterisks indicate the statistical significance between the original and permuted data (non-parametric test, p < 6.26e-249). **D**: Heat maps and violin plots of *k*-sim Jaccard for the selected TFs. In the heat maps, the rows and columns correspond to ChIP-seq samples and the color indicates *k*-sim Jaccard, whereas the color label indicates cell type classes. In the violin plots, the x-axis indicates the ChIP-seq sample pairs with the same and different cell type classes and the y-axis indicates *k*-sim Jaccard. Asterisks indicate the statistical significance between the ChIP-seq samples with the same and different cell type classes (Mann–Whitney U test, p < 0.05). **E**: Left: Pie chart of the ratio of cell type-dependent and non-cell type-dependent TFs. Right: schematic illustration of the cell type-dependent TFs. The TFs were excluded where those p-values were not calculated. **F**: Cell type-dependent TF similarity patterns by the weighted graph. Each node represents a TF, and the node in the center represents the query TF. The color of the edge indicates the *k*-sim Jaccard. For each query TF, the 15 TFs were selected whose difference in the *k*-sim Jaccard was the largest between the two cell type classes.

Next, we divided the MOCCS profiles for each TF by cell type class and compared two MOCCS profiles using the *k*-sim Jaccard because the *k*-sim Jaccard can explicitly quantify the overlaps of significant *k*-mers (**Materials and Methods**) (**Fig. 3A**). For example, JUN showed a high *k*-sim Jaccard in the same cell type class, which was statistically significant compared with different cell type classes (**Fig. 3D**). Similarly, we found cell type-dependent TFs such as FOS and GATA2 and non-cell type-dependent TFs such as MYC (**Fig. 3D**). Given these examples, we defined the TFs whose *k*-sim Jaccard showed statistical significance between the same and different cell type classes as cell type-dependent TFs (Mann–Whitney U test, p < 0.05). Overall, we found 42 cell type-dependent and 85 non-cell type-dependent TFs (33%) from 127 TFs. (**Fig. 3E, Figs. S6 and 7, Table S1**).

Additionally, we compared the TF similarity patterns of JUN and GATA2 between blood and pluripotent stem cells and identified different patterns among the cell type classes (**Fig. 3F**). In summary, the MOCCS profile comparisons revealed the cell type dependencies of TF-binding sequences and TF similarity patterns (**Fig. 3F**).

### Differentially recognized *k*-mers between two ChIP-seq samples

Given the sample-level differences in MOCCS profiles among ChIP-seq samples, we next focused on which *k*-mers showed different MOCCS2scores between two ChIP-seq samples. As differentially expressed genes in RNA-seq analysis, differential analysis of MOCCS2score would provide differential *k*-mers that are differentially recognized by TFs between two ChIP-seq samples (**Fig. 4A**). For each *k*-mer, we calculated the p-value of the difference in the MOCCS2score between two ChIP-seq samples and the corresponding q-value for multiple testing correction (**Materials and Methods**) and defined *k*-mers that satisfied q < 0.05 as differential *k*-mers. Using simulated datasets, we verified the q-values of differential *k*-mers (**Fig. S8A and B**), resulting in > 75% sensitivity and > 98% specificity, and controlled for the FDR (**Fig. 4B and C**) in five conditions.

**Figure 4.**
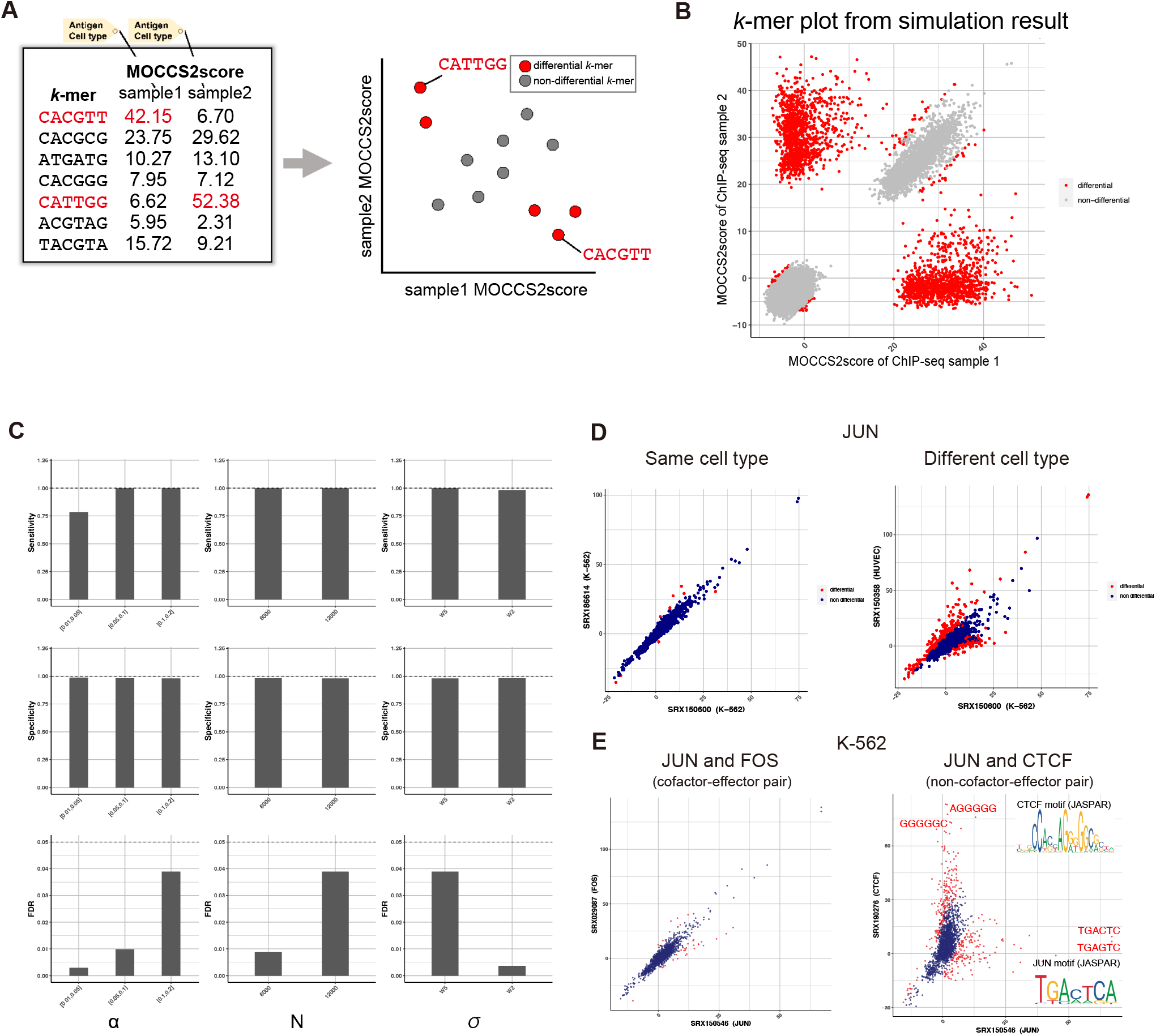
Differential analysis of *k*-mer profiles between ChIP-seq sample pairs can detect differentially recognized *k*-mers. **A**: Schematic overview of the simulation of the differential *k*-mer detection. **B**: Simulation results of the differential *k*-mer detection. The x- and y-axes represent the MOCCS2score of one ChIP-seq sample and a different ChIP-seq sample, respectively. Red and gray points represent differential *k*-mers (q < 0.05) and the other *k*-mers, respectively. **C**: Bar plots showing the sensitivity, specificity, and FDR of the detection of differential *k-*mers for different simulated conditions (**Fig. S8B**). α is the percentage of input sequences containing embedded “true significant *k*-mers”, N is the number of peaks in a ChIP-seq sample, and σ is the standard deviation of the embedded “true significant *k*-mers” from the center of the peak. **D**: Scatter plots of differential *k*-mers between two ChIP-seq samples with different cell types for the same TF (JUN). Red and blue points represent differential *k*-mers (q < 0.05) and the other *k*-mers, respectively. **E**: Scatter plots of differential *k*-mers between ChIP-seq sample pairs of different TFs in same cell types (K-562). Red and blue points represent differential *k*-mers (q < 0.05) and other *k*-mers, respectively. The PWM-supported differential *k*-mers and known PWM motifs (JASPAR) are shown in comparison between the JUN and CTCF ChIP-seq samples.

Differences in the biological contexts of ChIP-seq samples paralleled the number of differential *k*-mers between two ChIP-seq samples. For example, a comparison of the MOCCS profiles of JUN between the same cell types (K-562) and different cell types (K-562 and HUVEC) identified 10 (0.48%) and 293 (14.0%) differential *k*-mers, respectively (**Fig. 4D**). This indicates the detection of a higher number of differential *k*-mers in the comparison of ChIP-seq samples from different cell types than in the comparison of those from the same cell type. When we compared the MOCCS profiles between JUN and CTCF in the K-562 cell type (**Fig. 4E**), 293 differential *k*-mers (14.0%) were found, which was greater than the 38 differential *k*-mers (1.82%) found for the JUN and FOS pair (**Fig. 4E**). Because JUN dimerizes with FOS (34) but not with CTCF, the higher number of differential *k*-mers in the comparison of JUN and CTCF is reasonable. For JUN and CTCF, the differential *k*-mers with high MOCCS2scores contained PWM-supported *k*-mers (the *k*-mers with the top likelihoods from JASPAR) (**Fig. 4E**). Accordingly, differential *k*-mers can be applied to detect different TF-binding sequences between two different TFs. Thus, the differential *k*-mers in MOCCS profiles enable us to detect variable *k*-mers between two ChIP-seq samples.

### ΔMOCCS2score profiles agree with *in vitro* SNP-SELEX data and *in vivo* allele-specific binding data

We next hypothesized that the ΔMOCCS2score could indicate the single-nucleotide polymorphisms (SNPs) affecting TF binding. Specifically, the ΔMOCCS2score between two *k*-mers differing from each other by one nucleotide might reflect the effect of SNPs in the TF-binding region (**Fig. 5A**). Here, we use the ΔMOCCS2score as the difference in the MOCCS2score between two different *k*-mers in a single ChIP-seq sample.

**Figure 5.**
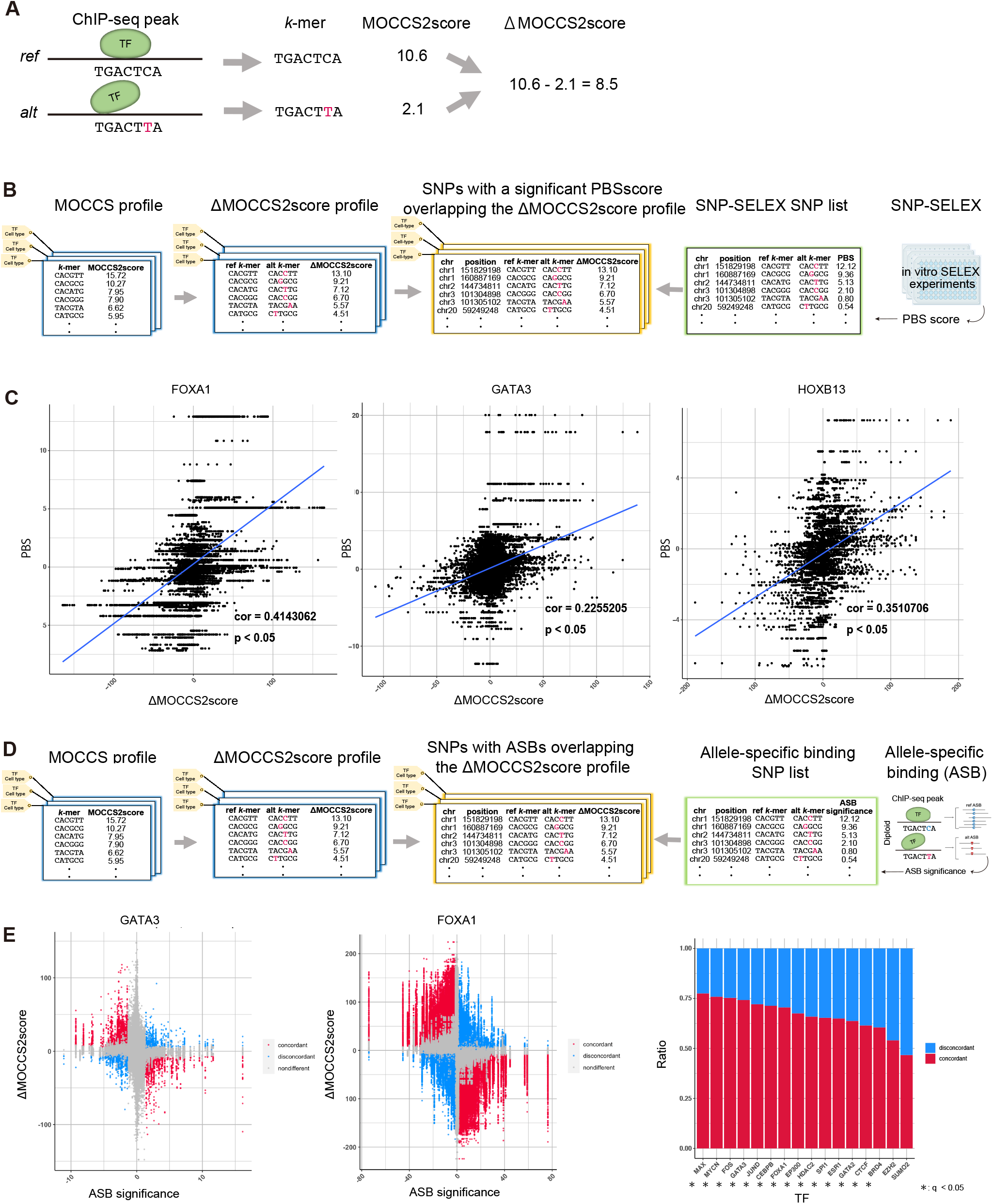
ΔMOCCS2score profiles agree with the *in vitro* SNP-SELEX data and *in vivo* ASB data. **A**: Schematic overview of the ΔMOCCS2score calculation for SNP-overlapping TF-binding *k*-mers. **B**: Data processing procedures to calculate the ΔMOCCS2score in SNP-overlapping TF-binding *k*-mers that exhibited significantly differential binding to at least one TF in SNP-SELEX experiments (35). **C**: Comparison between the PBS (SNP-SELEX) and ΔMOCCS2score. Each point represents a SNP corresponding to *k*-mer pairs (ref-*k*-mer and alt-*k*-mer). The Spearman correlation coefficient between the PBS and ΔMOCCS2score and the corresponding p-values (*t*-test) were calculated for each TF. The scatter plots contain ChIP-seq samples in all cell types of focal TFs. **D**: Data processing procedures to calculate the ΔMOCCS2score for *k*-mers overlapping SNPs with ASB events (36). **E**: Left and middle: Comparison between ASB significance and the ΔMOCCS2score. Each point represents a SNP corresponding to *k*-mer pairs (ref-*k*-mer and alt-*k*-mer). Red points are concordant SNPs. Right: Bar plots showing the ratios of concordant and discordant SNPs for each TF. Asterisks indicate a significant concordant ratio in the TFs (p-values were calculated from the empirical null distribution of the percentage of concordant SNPs, p < 0.05).

To verify whether the ΔMOCCS2score can evaluate the effect of SNPs overlapping TF-binding sequences, we compared the ΔMOCCS2score with the results of SNP-SELEX (35) (**Fig. 5B**). SNP-SELEX uses the preferential binding score (PBS) to assess the influence of SNPs that alter the *in vitro* binding specificity of TFs to DNA sequences. A positive and large PBS indicates stronger binding of the TFs, which is caused by changes from the reference allele to the alternative allele. We calculated the Spearman correlation coefficients between the PBS and ΔMOCCS2score for 10 TFs and found that 9 TFs had positive correlation coefficients (Student’s *t*-test, p < 0.05) (**Fig. 5C**). Furthermore, the SNPs located in the center of the *k*-mers showed stronger positive correlations between the PBS and ΔMOCCS2score (**Fig. S9A**), indicating that the ΔMOCCS2score correctly detected the effects of the SNPs. These results indicated that the ΔMOCCS2score was consistent with the *in vitro* SNP-SELEX findings.

Next, we compared the ΔMOCCS2score with allele-specific binding (ASB) significance (36), a measure of the *in vivo* ASB of SNPs based on ChIP-seq data (**Fig. 5D**). The ASB significance quantifies the influence of SNPs on binding affinity, and a negative and larger ASB significance indicates a stronger influence of TF-binding specificity caused by a change from the reference allele to the alternative allele. We evaluated the fraction of SNPs showing concordance between the ΔMOCCS2score and ASB significance. For example, 74% of the SNPs in GATA3 ChIP-seq data and 70% in FOXA1 ChIP-seq data were concordant (concordant SNPs) (**Fig. 5E, left**). Among the 16 tested TFs, 14 had significantly higher percentages of concordant SNPs compared with the experimental negative control (**Fig. 5E, right; Materials and Methods**), indicating that the ASB significance and ΔMOCCS2score were consistent. Furthermore, as with the consistency between the ASB significance and motif fold change from PWM motifs (36), the ΔMOCCS2score agreed with the motif fold change, indicating the consistency of the ΔMOCCS2score (**Fig. S9B**). Collectively, these results confirmed that the ΔMOCCS2score of two different *k*-mers could evaluate the effect of SNPs in TF-binding regions on the binding specificities of TFs.

### Evaluation of GWAS-SNPs in TF-binding regions and prediction of SNP-affected TFs through ΔMOCCS2score profiles

More than 90% of SNPs reported in genome-wide association studies (GWASs) are located in non-coding regions (37) and enriched in predicted transcriptional regulatory regions, called “cis-regulatory elements” (CREs) (38). However, it remains challenging to predict the SNPs affecting TF binding (39). Therefore, we used the ΔMOCCS2score to predict the TF binding for each GWAS-SNP (**Fig. 6A**).

**Figure 6.**
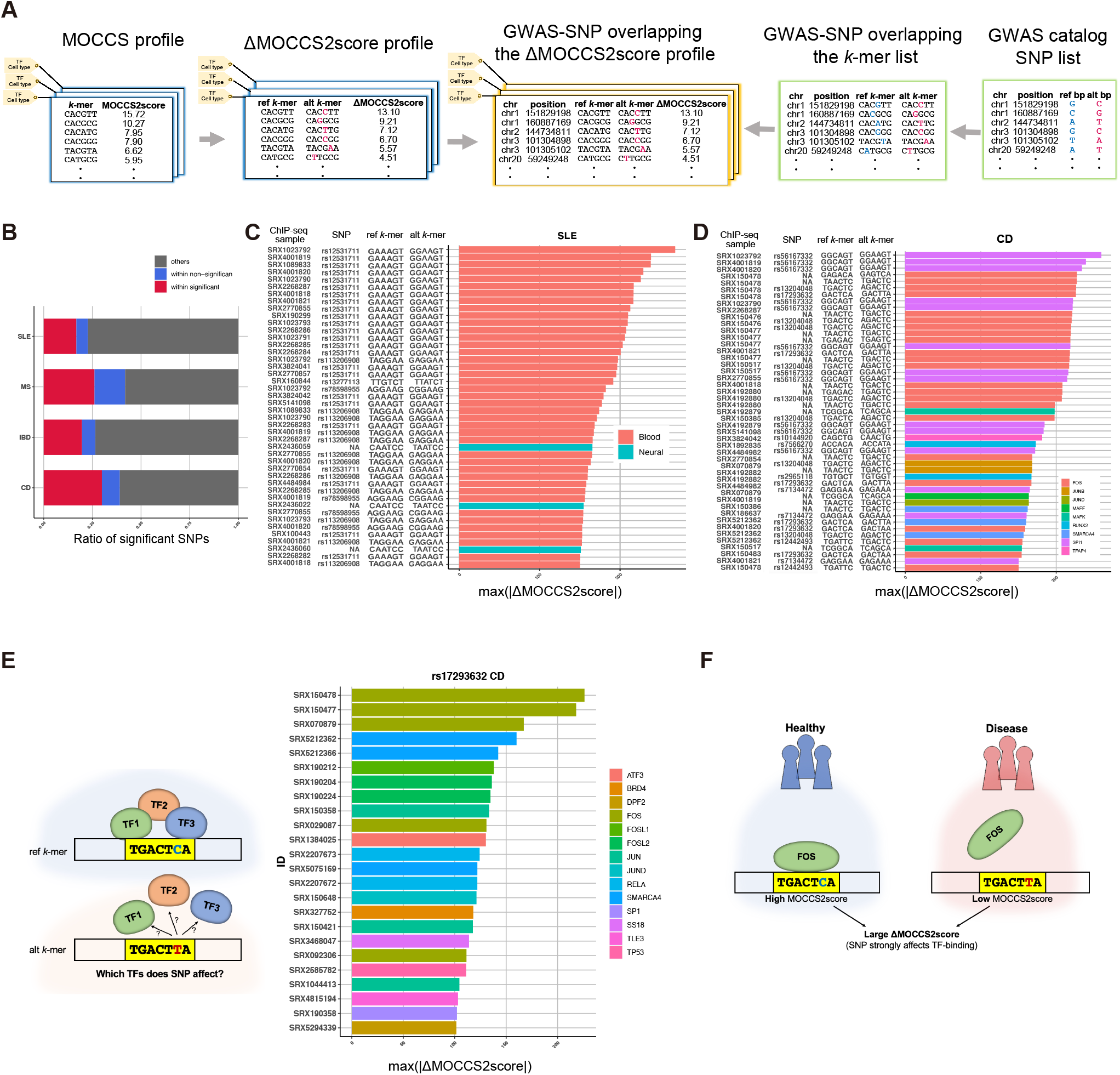
Evaluation of GWAS-SNPs in the TF-binding region and prediction of SNP-affected TFs through ΔMOCCS2score profiles. **A**: Data processing procedures to calculate the ΔMOCCS2score for *k*-mers overlapping GWAS-SNPs. **B**: Ratios of GWAS-SNPs divided by overlaps with ChIP-seq peaks and significant ΔMOCCS2scores (q < 0.05). The red fraction represents peak-overlapping SNPs with significant ΔMOCCS2scores and the blue fraction represents peak-overlapping SNPs with non-significant ΔMOCCS2scores. The gray fraction represents GWAS-SNPs outside of ChIP-seq peaks. **C**: ChIP-seq samples with high ΔMOCCS2scores (|ΔMOCCS2score| > 150) in each phenotype. Each SNP in a ChIP-seq sample has a ΔMOCCS2score. The bar graph shows SNPs rearranged by the |ΔMOCCS2score| and the ChIP-seq sample annotation (TF or cell type class). Each bar is each SNP from a ChIP-seq sample. **D**: (Left) Schema of the prediction of SNPs affecting TF binding. (Right) Prediction of SNPs affecting TF binding by ΔMOCCS2score profile (Crohn’s disease, rs17293632 C>T). Bar colors indicate TFs. The top three ChIP-seq samples with high absolute values of the ΔMOCCS2score were FOS. **E**: An example of GWAS-SNPs predicted to affect FOS binding by ΔMOCCS2score profiles in Crohn’s disease. Rs17293632 (C>T) might strongly affect the binding of FOS to the SNP-overlapping *k*-mer because it had the highest absolute value of the ΔMOCCS2score corresponding to the SNP-overlapping *k*-mer (**Fig. 6D**).

We selected four human disease phenotypes: systemic lupus erythematosus (SLE), multiple sclerosis (MS), inflammatory bowel disease (IBD), and Crohn’s disease (CD). Of the 626–971 GWAS-SNPs for each phenotype, 22.7%–45.8% overlapped the ChIP-seq peaks and 16.8%–29.8% showed a significant ΔMOCCS2score (q < 0.05) (**Fig. 6B**), suggesting that the ΔMOCCS2score could detect the effects of GWAS-SNPs. The GWAS-SNPs with significant ΔMOCCS2scores were distributed in each ChIP-seq sample (**Fig. S10A**) and for each TF (**Fig. S10B**). Given these trends, we next focused on the ChIP-seq samples in each disease and predicted the TFs and cell types whose binding specificity was influenced by SNPs.

SLE is an autoimmune disease that affects multiple organs, including the skin, joints, central nervous system, and kidneys (40). SLE GWAS-SNPs showed a high ΔMOCCS2score (|ΔMOCCS2score| > 150) when they overlapped the peaks in ChIP-seq samples of the blood cell type class, accounting for 74% of the top 100 SNPs (**Fig. 6C**). This result was consistent with previous reports showing the enrichment of overlapping SNPs for SLE in hematopoietic cells (41, 42). Moreover, SLE GWAS-SNPs were enriched with SPI1 ChIP-seq (**Fig. S11**) and the top SNPs for SLE corresponded to the known SPI1 motif. These results indicated that the ΔMOCCS2score could predict cell types in which the TF-binding specificity was possibly influenced by GWAS-SNPs.

For CD, ChIP-seq samples of SPI1 showed the highest |ΔMOCCS2score|, which accounted for 17% of the |ΔMOCCS2scores| of the top 100 SNPs, and FOS accounted for 28% of the |ΔMOCCS2scores| of the top 100 SNPs (**Fig. 6D**). The alt-*k*-mer of rs56167332 (GGAAGT) corresponded to the motif of SPI1, and the ref-*k*-mer of rs13204048 (TGACTC) corresponded to the motif of FOS, indicating that the predictive capability of the ΔMOCCS2score was reasonable. The enrichment of SPI1 and FOS was also evident in IBD GWAS-SNPs (**Fig. S11**), consistent with a previous report showing fold-enrichment of GWAS loci within regions marked by SPI1 binding (43). Among the |ΔMOCCS2scores| of the top 100 SNPs, we focused on rs17293632 (ref-*k*-mer: GACTCA; alt-*k*-mer: GACTTA), which has been reported to be a CD-associated variant and an ASB SNP for JUN and FOS (36, 44). Among the ChIP-seq samples whose peaks overlapped rs17293632 with a high |ΔMOCCS2score| (|ΔMOCCS2score| > 100), FOS had the top |ΔMOCCS2score| (|ΔMOCCS2score| = 226.1) and accounted for 40% of the |ΔMOCCS2scores| of the top 100 ChIP-seq samples (**Fig. 6E and F**), indicating that the ΔMOCCS2score could predict TFs whose binding specificity was influenced by GWAS-SNPs.

Finally, we confirmed that, as the allele frequency of GWAS-SNPs increased, the absolute values of the ΔMOCCS2score and the ratio of SNPs with a significant ΔMOCCS2score tended to decrease (**p < 0.001 using F-test; Fig. S12**). This is consistent with the fact that deleterious alleles tend to have lower allele frequencies in the human population (45). In summary, the ΔMOCCS2score obtained from the MOCCS profile could be applied to predict the combinations of TFs and cell types whose binding specificity is influenced by SNPs associated with human disease.

## Discussion

In this study, we proposed MOCCS profiles, the new representation of DNA-binding specificity of TFs for ChIP-seq samples, to capture features of TF-binding sequences. We used MOCCS2 to systematically calculate MOCCS profiles for >10,000 human TF ChIP-seq samples across diverse TFs and biological contexts. By comparing MOCCS profiles with conventional PWMs, we confirmed that these profiles can capture *k*-mers recognized by TFs. Moreover, by comparing the MOCCS profiles across ChIP-seq samples from different TFs and different cell types, we found that MOCCS profiles can capture TF-binding sequence similarities between (1) TFs of the same TF families and (2) cell types of the same cell type classes. We also found that one-third of TFs show cell type dependency in TF-binding sequences. Cell type-dependent TFs might pose challenges to the use of machine learning to predict TFBSs (46) and require more sophisticated methods, such as multi-task learning (47).

Given that the MOCCS profiles were representative of the ChIPed TF’s DNA binding specificities in ChIP-seq samples, we derived ΔMOCCS2scores, which reflect the differences in TF-binding specificity, based on the MOCCS profiles and used them to predict the effects of SNPs on TF binding. Using *in vitro* SNP-SELEX and *in vivo* ASB datasets, we confirmed that the ΔMOCCS2score analysis accurately predicted the SNPs affecting TF binding. Furthermore, we calculated the ΔMOCCS2scores for GWAS-SNPs of several diseases in the entire high-quality human ChIP-seq dataset and found candidate cell types and TFs associated with each disease. These results collectively show how the MOCCS profiles and ΔMOCCS2scores contribute to our understanding of TF-binding sequences.

In this study, we did not investigate the molecular bases of the cell type dependency of binding sequences. One possible mechanism is a change in chromatin accessibility and 3D chromatin structures, which have been associated with cell type-specific gene expression (48). This mechanism could be examined through a comparison of MOCCS profiles with chromatin accessibility and structures using DNase I-seq, ATAC-seq, and Hi-C data. Another possibility is that different TF partners alter the binding specificities, as systematically investigated using *in vitro* assays (49). We could address this possibility by comparing MOCCS profiles with TFBS colocalization patterns.

There are several possible future research directions. The first is to investigate the relationship between *k*-mer usage and other genomic features, including chromatin states, gene density, and gene functions. The second is to use the ΔMOCCS2score to interpret various types of mutation information, including mutation signatures (50) and indels (51). The third is to apply the ΔMOCCS2score analysis of GWAS-SNPs to drug target discovery by searching for SNPs affecting TF binding as well as candidate responsible cell types (52). The fourth is to investigate the diversity of TF-binding sequences among homologues of human TFs. For example, functional diversification of TF homologues parallels the diversification of MOCCS profiles in zebrafish (16). The fifth is to investigate interpositional dependencies within TF-binding motifs for all ChIP-seq data and the diversity of interpositional dependencies across cell types and TFs. Interpositional dependencies are not limited to directly adjacent nucleotides (53), and *k*-mer-base motif analyses have revealed interpositional dependencies in TF-binding motifs (25, 29). Finally, we could develop a userfriendly database to help researchers investigating gene expression regulations and human genetics. We are now developing a database to provide the precomputed results, including MOCCS profiles and ΔMOCCS2scores, for soft- and hard-filtered human ChIP-seq samples.

## Conclusions

Our results demonstrate the effectiveness of the MOCCS profile analysis for revealing the landscape of TF-binding sequences across diverse TFs and cell types in humans and that of ΔMOCCS2scores for interpreting GWAS-SNPs. We believe that the MOCCS profile analysis would provide the basis for investigating gene expression regulation and non-coding disease-associated variants in humans.

## Methods

### Preprocessing of ChIP-seq data

We obtained ChIP-seq peak data from ChIP-Atlas (https://chip-atlas.org/), trimmed each peak region in BED files to +/− 350 bp from the TFBSs (the centers of each peak region), and converted the BED files to FASTA files using the BEDTools getfasta tool for application to MOCCS2 with the options “--mask option, -low-count-threshold −1”. We obtained the annotations of TFs (antigen), cell type, and cell type class from the ChlP-Atlas.

### Sample filtering

To maintain the quality of the ChIP-seq samples and MOCCS profiles, we obtained quality metrics and set two kinds of thresholds, a soft filter and a hard filter.

#### Soft filter

For the soft filter, we obtained quality control metrics from processing logs (https://github.com/inutano/chip-atlas/wiki#tables-summarizing-metadata-and-files/) in ChlP-Atlas and read alignment rate from bowtie2 results. We set the thresholds as follows: number of mapped reads, 10,000,000; number of peaks, 100; and a mapping rate determined by mean - 2SD (54.09364).

#### Hard filter

For the hard filter, we used the quality metrics from DROMPAplus (54), a quality control tool for ChIP-seq experiments. To apply DROMPAplus, we obtained FASTQ files by using sra-tools_2.11.0.sif fasterq-dump or by downloading them from the DDBJ database, concatenating FASTQ files with the same SRX ID into one FASTQ file. We converted the FASTQ files to SAM files using bowtie2, converted the SAM files to BAM files using SAMtools, and applied DROMPAplus to the BAM files. We obtained 10,534 DROMPAplus output files, which contained five parameters for ChIP-seq quality control: library complexity, number of mapped reads, GC content, NSC (normalized strand cross-correlation coefficient), and Bu (background uniformity). We set the thresholds of the five parameters for the hard-filtered samples as follows: library complexity > 0.8; number of mapped reads >

10,000,000; GC content < 60; NSC > 2.0; Bu > 0.8; and number of peaks > 100. In addition, we removed ChIP-seq samples of GFP, epitope tags, BrdU, and biotin and finally selected 2,976 samples in the hard filter.

### Calculation of the MOCCS2score by MOCCS2

MOCCS2 clarifies TF-binding *k*-mers from ChIP-seq peak calling data (23, 29). Specifically, MOCCS2 quantifies the sharpness of the histogram as it appears for each *k*-mer around TFBSs as the AUC score, and then calculates the MOCCS2score for each *k*-mer by normalizing the AUC scores. The AUC score is the area under the cumulative relative frequency curve of the appearance of each *k*-mer sequence against the distance from the TFBSs. *W* is the size of the analyzed window where *k*-mer sequences are sought at around ChIP-peak positions and *n* is the number of *k*-mer appearances in ChIP-seq samples. Let *f*(*x*) be the appearance count of each *k*-mer sequence at the position ±*x* bp (*x* ∈ [1, *W*]) away from the TFBSs, then the cumulative relative frequency distribution *F*(*x*) for the *k*-mer sequence is calculated as follows:

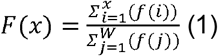

and its AUC score is calculated as follows:

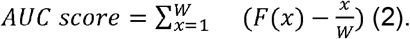

Note that the AUC score becomes larger as the shape becomes sharper.

MOCCS2score of each *k*-mer was defined as a relative value of the AUC score normalized by the SD at its appearance count. Some irrelevant *k*-mers with low appearance counts show high AUC scores due to large standard deviations (SDs) of the AUC of the irrelevant sequences (29). To compensate for the falsely high values of the AUC scores, we defined a MOCCS2score for each *k*-mer as the AUC score normalized by the SD at its appearance count. The SDs of the AUC scores is calculated as 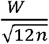, as derived in (29).

Then, the MOCCS2score is calculated as follows:

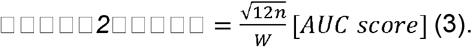

### p-value of the MOCCS2score by MOCCS2

When a *k*-mer randomly appears within ±*W* bp from a TFBS, *f*(*x*) (distribution of the *k*-mer position from the TFBS) follows the uniform distribution *U*(0, *W*) (23, 29). Then, the AUC score is regarded as the sample mean of the uniform distribution *U*(0, *W*) subtracted by 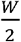:

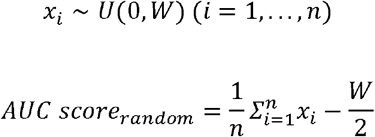

According to the central limit theorem, when *n* is sufficiently large, the sample mean of *U*(0, *W*) approximately follows the normal distribution 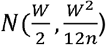 because the mean and variance of *U*(0, *W*) are 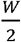 and 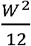, respectively. As the AUC score is the sample mean of *U*(0, *W*) subtracted by 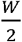 when *n* is sufficiently large, the AUC score approximately follows the nominal distribution: 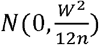. Then, the p-value of the observed AUC score *s* is calculated as follows:

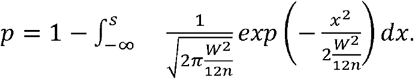

In this study, we defined the p-value of the MOCCS2score of a *k*-mer as the p-value of the corresponding AUC score.

We evaluated the p-value of the AUC score by comparing it with empirical p-values based on a simulation experiment. Then, each *k*-mer position relative to the peak center was simulated by sampling from *U*(0, *W*). The AUC score was calculated as sampled_means - W/2. We set *n* = 100 (assumed to be a *k*-mer with 100 counts) and *W* = 250. The simulation was repeated 10,000 times and the empirical distribution of the AUC score was obtained.

The ratio of the empirical standard deviation to the theoretical value was 1.0064, indicating that the p-value based on the central limit theorem and the p-value calculated from the empirical cumulative distribution based on the simulation results were roughly consistent.

### Significant *k*-mer detection

The p-values of the MOCCS2scores of *k*-mers were calculated in each sample. The q-value was calculated from the p-values from each sample by the “p.adjust” function in the “stats” package in R for multiple testing correction. *k*-mers with q < 0.05 were considered significant *k*-mers.

### Evaluation of the prediction performance of PWM-supported *k*-mers based on the MOCCS2score

#### Calculation of the likelihood for each *k*-mer by the PWM motif

We downloaded the motif PWMs from HOCOMOCO

(https://hocomoco11.autosome.ru/downloads_v11). For each PWM, the likelihood was calculated for each *k*-mer by multiplication of each base probability across positions in the PWM. We calculated the likelihood by shifting a *k*-mer with a PWM for all combinations and then used the maximum determined value.

#### Evaluation of the performance by the MOCCS2score to detect PWM-supported *k*-mers

We defined *k*-mers with a high likelihood (top 10%) as “positive”*k*-mers (PWM-supported *k*-mers) and the other *k*-mers as “negative”*k*-mers. We predicted the classification performance of the “positive”*k*-mers using the MOCCS2score of ChIP-seq samples of the focal TF and calculated the area under the ROC (AUROC).

### MOCCS profile comparison

#### Calculation of the *k*-sim Pearson and Jaccard and the peak overlap index

Between each MOCCS profile pair that passed the hard filter, we defined two similarity indices: the *k*-sim Pearson and *k*-sim Jaccard (**Fig. 2A**).The *k*-sim Pearson of a pair of MOCCS profiles was defined as the Pearson correlation coefficient of the MOCCS profiles after setting the MOCCS2scores of non-significant *k*-mers to 0. Note that we excluded MOCCS profiles whose all *k*-mer were non-significant from **Figures 2 and 3** and **S4, S5, S6, and S7**. The *k*-sim Jaccard of a pair of MOCCS profiles was defined as the Jaccard index of significant *k*-mers in two MOCCS profiles (q < 0.05), A and B, as follows:

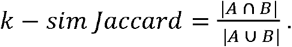

Note that the *k*-sim Pearson calculates the similarity, together with the sign of correlation, by considering the value of each significant *k*-mer, whereas the *k*-sim Jaccard calculates the overlap degrees of significant *k*-mers.

Based on the BED files obtained from ChIP-Atlas (21), we also calculated the peak overlap index as a control, which directly reflects the degree of the peak overlap regions, in the three steps. First, for a pair of ChIP-seq samples (indexed with 1 and 2), we calculated *n1all* and *n2all* as the total numbers of peaks in ChIP-seq samples 1 and 2, respectively. Second, we counted *n1* (*n2*) as the number of peaks in ChIP-seq sample 1 (2) that overlapped peaks in ChIP-seq sample 2 (1) using BEDTools with the intersect option (intersect -u -a -b) (55). Third, we calculated the peak overlap index as follows:

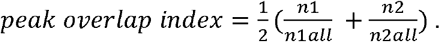

To validate the *k*-sim Pearson and Jaccard, we compared the *k-sim* Pearson (**Fig. 2B**) and Jaccard (**Fig. S4**) with the peak overlap index. Note that we excluded CTCF from **Figures 2** and **S4 and S5**. In the grouping of the MOCCS profile pairs, we excluded the pairs in which either sample annotation included “Unclassified”, “Others”, or “No annotation”.

#### UMAP visualization of MOCCS profiles and statistical tests

We performed UMAP on the set of MOCCS profiles using the R package “umap” (56). For the parameters of the package, we set the metric as “pearson” and the spread as 10. We colored these ChIP-seq samples on the UMAP plot by TF, TF family, or cell type class.

We performed the permutation test on the same annotation ratio of the TF, TF family, or cell type class for the top three neighboring ChIP-seq sample pairs defined by the *k*-sim Pearson. We conducted 1,000 permutations and collected the mean value of the same annotation ratio for the top three neighboring ChIP-seq sample pairs in each permutation trial (**Figure 2D and 3C)**.

We performed chi-square tests on TF, TF family, and cell type class (**Fig. S5**). In the chi-square test, we summed the same or different annotation ratios in the top three or not top three neighboring sample pairs defined by *k*-sim Pearson, respectively. Note that, we excluded CTCF from these UMAP procedures (**Fig. 2C and D, Fig. 3B and C**). We excluded unknown pairs whose annotations included “Unclassified”, “Others”, or “No annotation”.

#### Evaluation of TF similarity patterns by the *k*-sim Pearson

We calculated the *k*-sim Pearson among different types of TFs in a cell type class. We selected JUN in blood, FOS in blood, FOXF1 in digestive tract, and ELK1 in uterus as the query TFs and calculated the *k*-sim Pearson among the TFs whose ChIP-seq samples passed the hard filter. We extracted the TFs whose *k*-sim Pearson values were in the top ten for each TF queried, which were visualized as bipartite graphs.

#### Evaluation of cell type-dependent TFs by the *k*-sim Jaccard

We calculated the *k*-sim Jaccard for all pairs of MOCCS profiles in each TF. We visualized these *k*-sim Jaccard values as heat maps grouped by cell type class. We also visualized these *k*-sim Jaccard values as violin plots by dividing them into the same or different cell type classes. We investigated the statistical significance of the *k*-sim Jaccard between these same and different cell type class groups by using the Mann–Whitney U test. We denoted TFs whose *k*-sim Jaccard showed statistical significance between the same and different cell type classes as cell type-dependent TFs. In the calculation of the ratio of cell type-dependent and non-cell type-dependent TFs, we excluded TFs whose p-values were not calculated.

Among the cell type-dependent TFs, we selected JUN and GATA2 as the query TFs to compare TF similarity patterns between the two cell type classes. We selected the 15 TFs with the largest difference in the *k*-sim Jaccard between the two cell type classes. We selected the two cell type classes where the available TFs were the top two.

### Differential *k*-mer detection between ChIP-seq samples

#### Algorithm

To detect *k*-mers that are differentially recognized between two samples, we determine differential *k*-mers as follows. *W* is the size of the search window of *k*-mer occurrences around TFBSs. *n_i_* and *n_j_* are the number of appearances of *k*-mers *i* and *j*, respectively. When *k*-mers *i* and *j* appear randomly around TFBSs, the AUC scores of *i* and *j* follow the normal distributions *N*(0, *W*^2^/12*n_i_*) and *N*(0, *W*^2^/12*n_j_*), respectively. Then, the difference in the AUC scores between two *k*-mers can be regarded as the difference in the means between two normally distributed populations with unequal variance. In such a case, we can apply a two-sample z-test (57), which tests the hypothesis that two normally distributed populations with unequal variance have equal means. Let □^2^_*i*_ and □^2^_*j*_ be the variance of each *k*-mer distribution. When we assume that the variance of the AUC score is constant regardless of the value of the AUC score, the test statistics are:

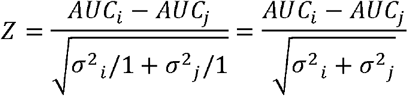

and follow the standard normal distribution. Then, the difference in the AUC score of the two *k*-mers *AUC_i_* – *AUC_j_* follows the normal distribution 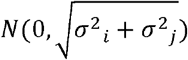. This approach is also applied to the statistical testing of the ΔAUC score (difference in AUC scores between two samples) and the p-values are calculated from the normal distribution 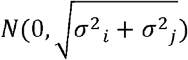.

#### Simulation of differential *k*-mer detection

To validate the differential *k*-mer detection method, we prepared simulation ChIP-seq peak data. We generated two random ChIP-seq samples (S1 and S2) with *N* peaks and comprising random sequences with length 2*W* + 1. Predetermined *k*-mers were embedded in the random sequences as follows:

- A: significant *k*-mers in S1 or S2 and non-differential *k*-mers
- B: significant *k*-mers in S1 or S2 and differential *k*-mers
  - B1: significant *k*-mers in S1 or S2 and differential *k*-mers in the S1 condition
  - B2: significant *k*-mers in S1 or S2 and differential *k*-mers in the S2 condition
- C: non-significant *k*-mers in S1 or S2 and non-differential *k*-mers

After each *k*-mer was embedded, each ChIP-seq sample (S1 and S2) was applied to MOCCS2 and the *p*-value of the difference in the MOCCS2score was calculated.

We set *l* (number of embedded differential *k*-mers) and *m* (number of embedded significant *k*-mers) as 45 and 90 based on the number of differential and significant *k*-mers in real ChIP-seq samples, respectively. We used *l* as the number of differential *k*-mers, which excluded 1-bp- or 2-bp-shifted *k*-mers from the original number of differential *k*-mers (200) in real ChIP-seq samples because we embedded *k*-mers without 1-bp- or 2-bp-shifted *k*-mers in the simulated ChIP-seq samples.

### Calculation of the ΔMOCCS2score in a single ChIP-seq sample and comparison of the ΔMOCCS2score with the SNP-SELEX, ASB SNPs, and GWAS-SNPs

#### Preparation of SNP-overlapping *k*-mer lists and the ΔMOCCS2score profile

To test whether the ΔMOCCS2score is consistent for assessing SNP-overlapping TF-binding regions, we used SNP-SELEX experiments (35), ASB SNP data (36), and SNP lists in the GWAS catalog (58) (https://www.ebi.ac.uk/gwas/). We obtained the *k*-mers overlapping SNPs from the reference genome (hg38). We defined *k*-mers obtained from the reference genome as ref*-k-*mers and defined *k*-mers that replaced the reference allele with the alternative allele as alt*-k*-mers, creating pairs of ref-*k*-mers and alt-*k*-mers. There are *k* possible positions for SNPs in the *k*-mer, so that the positions of SNPs in the *k*-mer were shifted from the 1st to the *k*th position from the left of the *k*-mer, creating *k* different ref-*k*-mers for each *k*-mer.

From the *k*-mer list, we obtained the AUC score, count, and MOCCS2score corresponding to both the ref-*k*-mer and alt-*k*-mer. We calculated the ΔMOCCS2score and its q-value as in the differential *k*-mer detection algorithm and generated the ΔMOCCS2score profile for each ChIP-seq sample.

#### Comparison of the ΔMOCCS2score with the SNP-SELEX results

We obtained the SNP-SELEX results from GSE118725(35) and lifted them over from hg19 to hg38. We selected the SNP list overlapping the ChIP-seq sample peak region and obtained the *k*-mer overlapping the SNP from the reference genome (hg38) (ref-*k*-mer). We created ref-*k*-mer and alt-*k*-mer pairs, obtained the AUC score, count, and MOCCS2score for each *k*-mer from the MOCCS profile, and calculated the ΔMOCCS2score, p-value, and q-value for each ChIP-seq sample. The SELEX results contain the PBS (preferential binding score) for each SNP. We calculated the correlation coefficient (Spearman) between the ΔMOCCS2score and PBS for each ChIP-seq sample of the same TF.

#### Comparison of the ΔMOCCS2score with the ASB SNPs

We obtained SNP lists from the ADASTRA database (36) (https://adastra.autosome.ru/zanthar, Release Susan v3.5.2), which contains ASB events and the corresponding ASB significance across 674 TFs and 337 cell types. The ASB significance indicates the changes in TF-binding specificity induced by ASB SNPs. We selected SNPs overlapping the peak regions from the ChIP-seq samples and obtained the *k*-mer overlapping the SNP from the reference genome (hg38) (ref-*k*-mer). After obtaining the alt*-k-*mer corresponding to the ref*-k-*mer, we determined the AUC score, count, and MOCCS2score from the MOCCS profile and calculated the ΔMOCCS2score, p-value, and q-value corresponding to the *k*-mer pair (ref*-k*-mer and alt*-k*-mer). A negative and larger ASB significance indicates stronger influence of TF-binding specificity caused by the change from the reference allele to the alternative allele. We defined concordant SNPs between the ΔMOCCS2score and ASB significance as those satisfying the following conditions: (1) ΔMOCCS2score is significant (q < 0.05); (2) |ASB significance| is significant (FDR < 0.05); and (3) the direction of change induced by the SNP is the same between the ΔMOCCS2score and ASB significance. Based on this definition, we calculated the ratio of concordant SNPs and discordant SNPs.

For each TF, we performed a permutation test on the percentage of concordant SNPs by shuffling ΔMOCCS2score profiles and calculated the p-value. We conducted 100 permutations. Furthermore, we obtained the fold change of the PWM from the ADASTRA database and calculated the Spearman’s correlation between the PWM motif fold change and the ΔMOCCS2score.

#### Evaluation of the ΔMOCCS2scores of GWAS-SNPs

We obtained GWAS-SNP data from the GWAS catalog (https://www.ebi.ac.uk/gwas/) for the following phenotypes: IBD (EFO_0003767), CD (EFO_0000384), MS (EFO_0003885), and SLE (EFO_0002690).

After selecting the SNPs overlapping the peaks of the ChIP-seq samples, we obtained the *k*-mer overlapping the SNPs from the reference genome (hg38) (ref-*k*-mer). After obtaining the alt-*k*-mer by changing one nucleotide of the ref-*k*-mer, we calculated the ΔMOCCS2score with a p-value and q-value for each ref*-k*-mer and alt-*k*-mer pair.

The association between the allele frequency and absolute values of the ΔMOCCS2score or ratio of SNPs with a significant ΔMOCCS2score was tested using linear regression. We set (1) allele frequency or (2) rank of allele frequency after their categorization into five bins as an explanatory variable, and (1) the absolute values of the ΔMOCCS2score or (2) the fraction of SNPs with a significant ΔMOCCS2score as a response. The p-values of regression coefficients were calculated using F-test.

## Abbreviations

ASB: Allele-specific binding
AUROC: Area Under Receiver Operating Characteristic Curve
ChIP-seq: Chromatin immunoprecipitation sequencing
GWAS: Genome-wide association study
PBS: Preferential binding score
PWM: Position weight matrix
SNP: Single-nucleotide polymorphism
TF: Transcription factor
TFBS: Transcription factor binding site
MOCCS: Motif centrality analysis of ChIP-seq

## Declarations

### Ethics approval and consent to participate

Not applicable.

### Consent for publication

All authors have approved the manuscript for submission.

### Availability of data and materials

The codes and data are available at the GitHub repository (https://github.com/bioinfo-tsukuba/MOCCS_paper_public.git). The following files are available at the figshare repository (10.6084/m9.figshare.19333646): (1) SNPs with a significant PBSscore overlapping the ΔMOCCS2score profile (SELEX_dMOCCS2score.tsv); (2) SNPs with ASBs overlapping the ΔMOCCS2score profile (ASB_dMOCCS2score.tsv); (3) GWAS-SNP overlapping the ΔMOCCS2score profile (GWAS_dMOCCS2score.tsv).

### Authors’ contributions

S.T., T.T., and H.O. conceived the project. S.T. and T.T. collected data. S.T., T.T., and H.O. performed analysis. H.M. devised methods for calculating the p-values of the MOCCS2score and ΔMOCCS2score. S.T., T.T., and H.O. wrote the manuscript. T.T. and H.O. conducted project administration. H.O. supervised the project. All authors discussed the results and approved the manuscript.

## Acknowledgements

Computations were partially performed on the National Institute of Genetics (NIG) supercomputer at the Research Organization of Information and Systems (ROIS) National Institute of Genetics, Japan. We thank Tazro Ohta (Database Center for Life Science, ROIS) and Dr. Shinya Oki (Kyoto University) for technical assistance on the use of the ChIP-Atlas data. We thank Dr. Y Tanizawa (NIG) and Dr. Manabu Ishii (Workflow meetup) for technical advice on NIG supercomputer systems. We also thank Dr. Ryuichiro Nakato (The University of Tokyo) for technical advice on the parameters of DROMPAplus.

## Figures, tables, and additional files

Figure S1 Filtering of ChIP-seq samples

**A**: Schematic overview of ChIP-seq sample filtering.

**B**: Distribution of each quality control metric of ChIP-seq sample filtering for samples that passed the hard filter (pink) and the others (blue).

Figure S2 Simulation of significant *k*-mer detection

**A**: The procedure for generating simulated datasets. We simulated data by embedding a “true significant *k*-mer” within random sequences, applying MOCCS2, and calculating the q-values of the MOCCS2score for each *k*-mer.

**B**: Parameters in each simulation condition from #1 to #5. α is the percentage of input sequences containing embedded “true significant *k*-mers”, N is the number of peaks in a ChIP-seq sample, and σ is the standard deviation of the embedded “true significant *k-*mers” from the center of the peak.

**C**: Simulation results for significant *k*-mer detection. The top row is sensitivity, the second row is specificity, and the bottom row is the FDR for detecting “true significant *k*-mers”.

Figure S3 Number of peaks and significant *k*-mers in MOCCS profiles

**A**: The number of peaks in MOCCS profiles. The x-axis represents the log-transformed number of peaks with a base of 10 and the y-axis represents the number of ChIP-seq samples.

**B**: Relationship between the number of peaks and significant *k-*mers in MOCCS profiles (left, q < 0.05; right, q < 0.01).

Figure S4 Comparisons of the *k*-sim Pearson and Jaccard and peak overlap indices

**A**: Comparisons of the *k*-sim Jaccard and Pearson and peak overlap indices.

**B**: Two-dimensional density plot of *k*-sim Jaccard or Pearson with the peak overlap index.

**C**: Correlation coefficient of the *k*-sim Jaccard or Pearson with the peak overlap index in each group. The y-axis indicates the Spearman’s correlation coefficient. The red and blue colors indicate the *k*-sim Pearson and Jaccard, respectively.

Figure S5 Chi-square testing of TF, TF family, and cell type class

**A**: Chi-square testing of TF. The color of the heat map indicates the summed ratio in each group. We summed the same or different annotation ratios in the top three or not top three neighboring sample pairs defined by the *k*-sim Pearson, respectively. The color label indicates each group.

**B** and **C**: Chi-square testing of TF family and cell type class. The heatmap color and the color label indicate the summed ratios and groups similar to A, respectively.

Figure S6 Heat maps of all of the cell type-dependent TFs

The heat map color indicates the *k*-sim Jaccard. The color label of the heat maps indicates the cell type classes. Asterisks indicate statistical significance of ChIP-seq samples with the same and different cell type classes (Mann–Whitney U test, p < 0.05).

Figure S7 Violin plots of all cell type-dependent TFs

The y-axis indicates the *k*-sim Jaccard. The same and different groups are arranged along the x-axis. Asterisks indicate the statistical significance of the ChIP-seq samples with the same and different cell type classes (Mann–Whitney U test, p < 0.05).

Figure S8 Simulation of differential *k*-mer detection

**A**: Simulated data processing. We prepared simulated data with an embedded “true differential *k*-mer” and “true significant *k*-mer” by embedding a “true”*k*-mer within α% of a randomly generated sample of 2W +1 - bp (W = 350) DNA sequences and applying

MOCCS2. “True significant *k*-mers” were embedded following a normal distribution whose mean was W + 1 and whose standard deviation was σ. “True differential *k*-mers” were embedded in S1 (or S2), the same as “true significant *k*-mers”, and were embedded in S2

(or S1) following a uniform distribution whose mean was 1 and whose standard deviation was (2 × W + 1) - (k - 1).

**B**: Parameters in each simulation condition from #1 to #5. *L* is the number of differential *k*-mers and *m* is the number of significant *k*-mers.

Figure S9 ΔMOCCS2score profiles agree with the *in vitro SNP-SELEX* and PWM motif fold change

**A**: Difference in the ΔMOCCS2score profile consistency among the positions of SNPs in *k*-mers. The kth SNP position means the kth allele from the left in the *k*-mer.

**B**: The ΔMOCCS2score agrees with the PWM motif fold change.

Figure S10 Number of differential peak-overlapping GWAS-SNPs

**A**: Number of peak-overlapping GWAS-SNPs with significant ΔMOCCS2scores of Crohn’s disease in one ChIP-seq sample. The red fraction shows the number of GWAS-SNPs with significant ΔMOCCS2scores (q < 0.05).

**B**: Number of differential peak-overlapping GWAS-SNPs of Crohn’s disease in the ChIP-seq samples of the same TFs.

Figure S11 Prediction of SNP-affected TFs and cell type classes through ΔMOCCS2score profiles

Top ChIP-seq samples with high ΔMOCCS2scores in each phenotype (IBD, inflammatory bowel disease; CD, Crohn’s disease; MS, multiple sclerosis; SLE, systemic lupus erythematosus). Each SNP in a ChIP-seq sample has a ΔMOCCS2score, and the SNPs have been arranged by the ΔMOCCS2score. Each bar is each SNP from a ChIP-seq sample. Bar graph colors show TFs or cell type class.

Figure S12 Association between the allele frequency and

ΔMOCCS2score

**A**: Association between the allele frequency and the absolute values of the ΔMOCCS2score.

**B**: Association between the allele frequency and the ratio of SNPs with a significant ΔMOCCS2score.

Table S1 List of cell type-dependent TFs

