## Supplementary Figures for "MOCCS profile analysis clarifies the cell type dependency of transcription factor-binding sequences and cis-regulatory SNPs in humans"

A

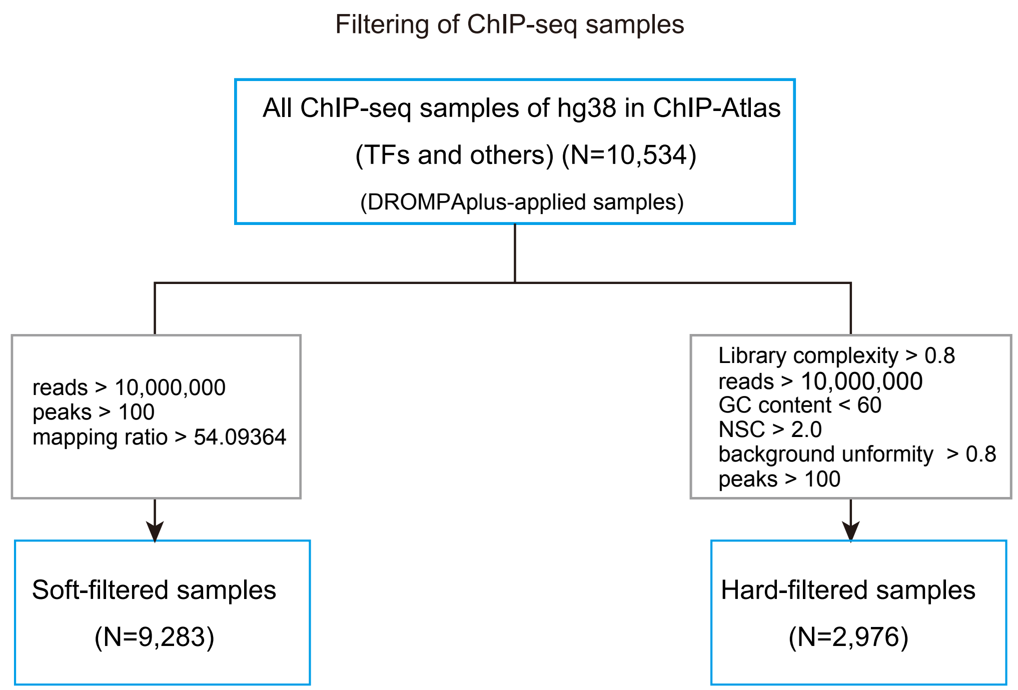


B

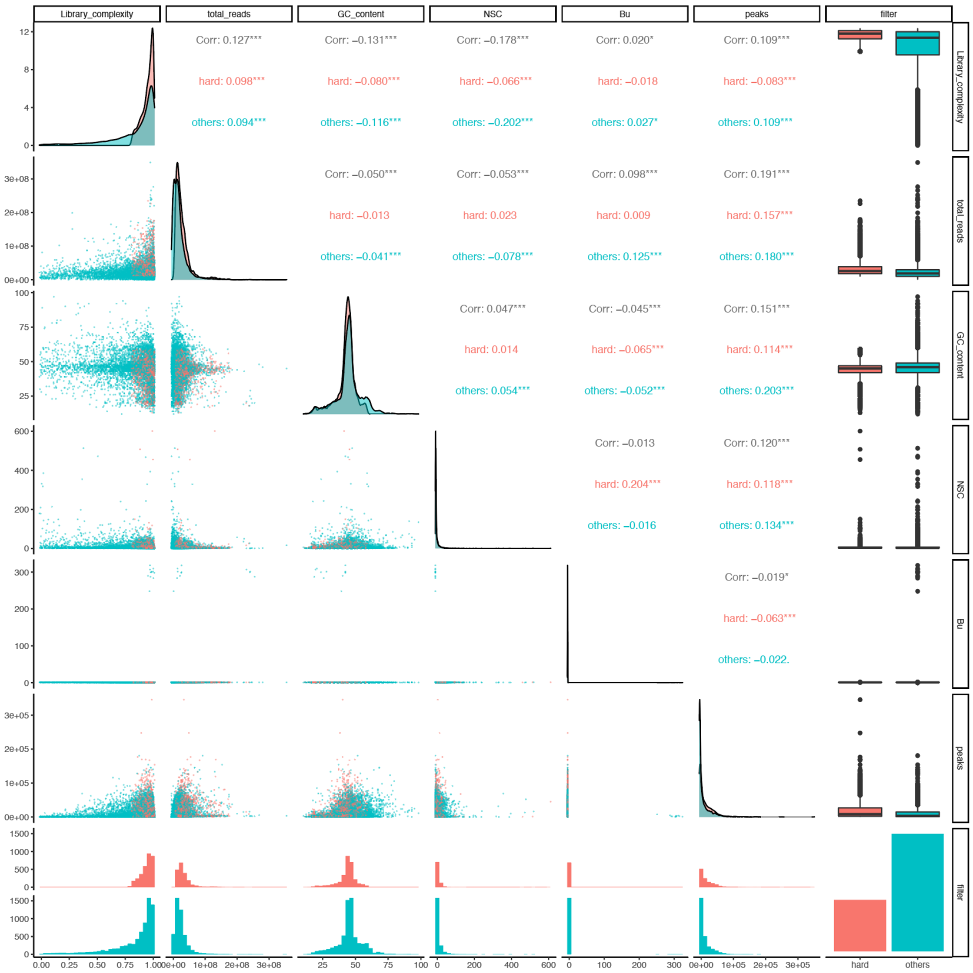


Figure S1. Filtering of ChIP-seq samples. A. Schematic overview of ChIP-seq sample filterings. B. Distribution of each quality control metric of ChIP-seq sample filtering for samples that passed the hard filter (pink) and the others (blue).

Tahara *et al.*, Figure S1

A

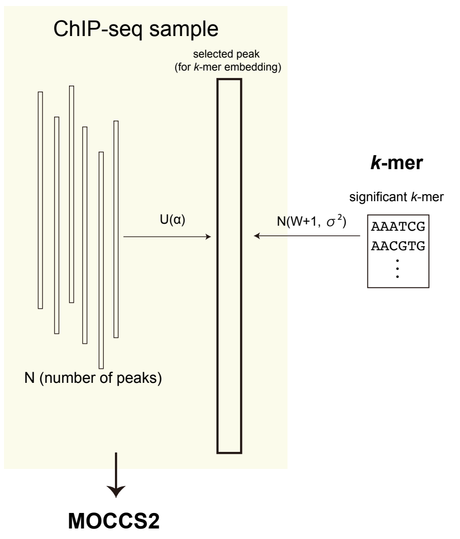


B

Parameters for simulation (significant *k*-mer)

| simulation | α | N | σ |
|---|---|---|---|
| #1 | [0.1, 0.2] | 12,000 | W/5 |
| #2 | [0.01, 0.05] | 12,000 | W/5 |
| #3 | [0.05, 0.1] | 12,000 | W/5 |
| #4 | [0.1, 0.2] | 6,000 | W/5 |
| #5 | [0.1, 0.2] | 12,000 | W/2 |

C

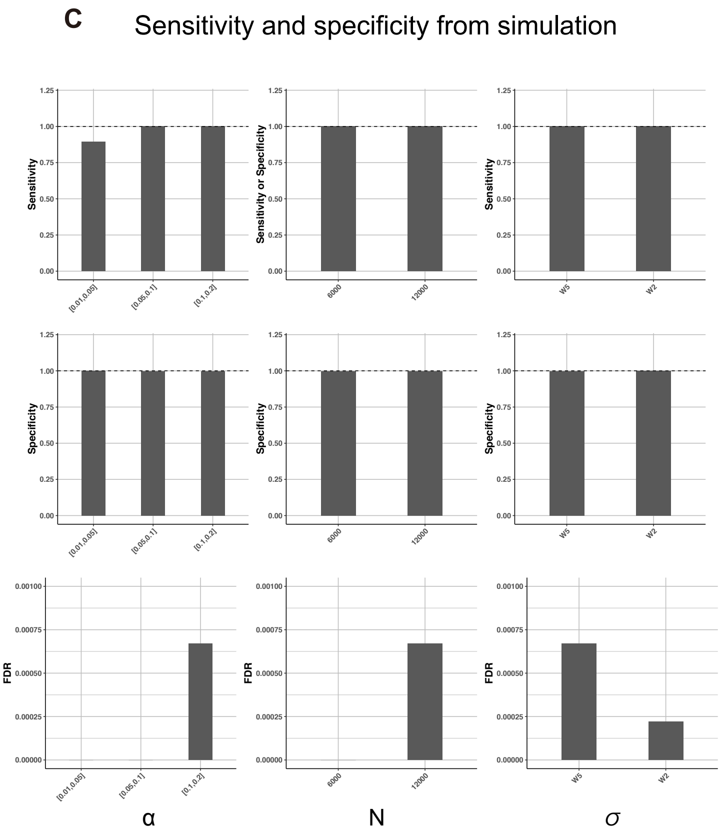


Figure S2. Simulation of significant *k*-mer detection. A. The procedure for generating simulated datasets. We simulated data by embedding a “true significant k-mer” within random sequences, applying MOCCS2, and calculating the q-values of the MOCCS2score for each k-mer. B. Parameters in each simulation condition from #1 to #5. α is the percentage of input sequences containing embedded “true significant k-mers” , N is the number of peaks in a ChIP-seq sample, and σ is the standard deviation of the embedded “true significant k-mers” from the center of the peak. C. Simulation results for significant k-mer detection. The top row is sensitivity, the second row is specificity, and the bottom row is the FDR for detecting “true significant k-mers” .

Number of peaks and significant *k*-mers in MOCCS profiles

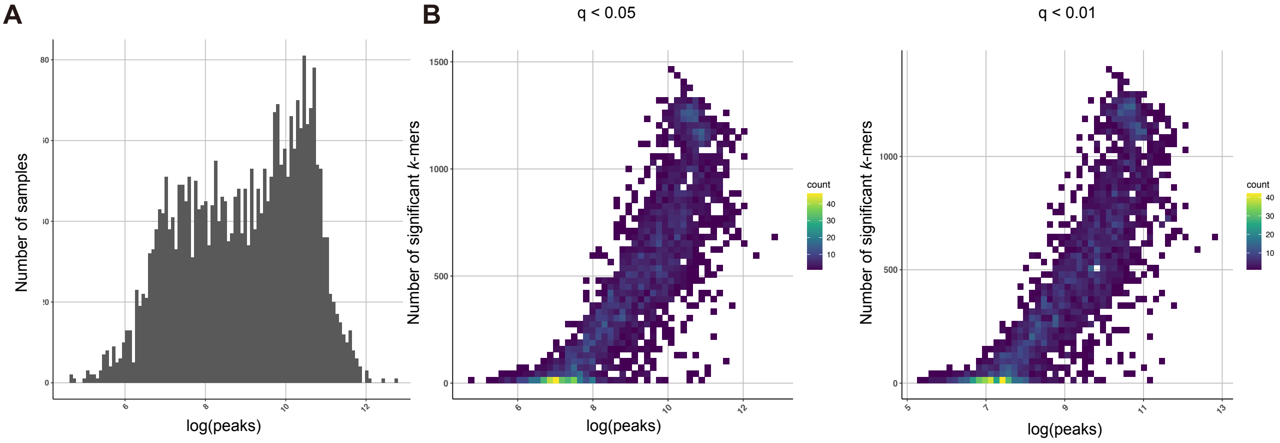


Figure S3. Number of peaks and significant *k*-mers in MOCCS profiles. A. The number of peaks in MOCCS profiles. The x-axis represents the log-transformed number of peaks with a base of 10 and the y-axis represents the number of ChIP-seq samples. B. Relationship between the number of peaks and significant *k*-mers in MOCCS profiles (left, $q < 0.05$; right, $q < 0.01$).

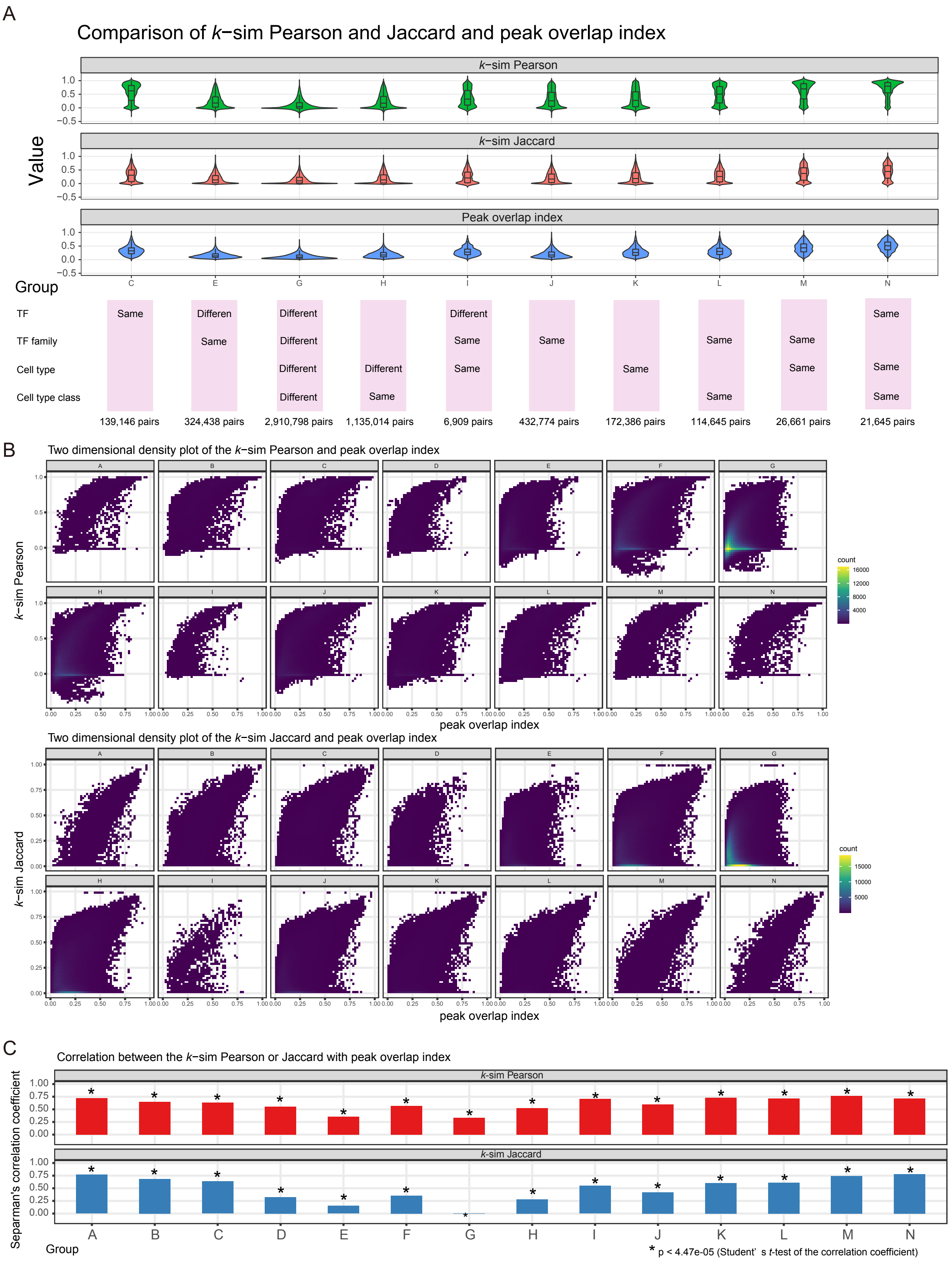


Figure S4. Comparisons of the *k*-sim Pearson and Jaccard and peak overlap indices. A. Comparisons of the *k*-sim Jaccard and Pearson and peak overlap indices. B. Two-dimensional density plot of *k*-sim Jaccard or Pearson with the peak overlap index. C. Correlation coefficient of the *k*-sim Jaccard or Pearson with the peak overlap index in each group. The y-axis indicates the Spearman' s correlation coefficient. The red and blue colors indicate the *k*-sim Pearson and Jaccard, respectively.

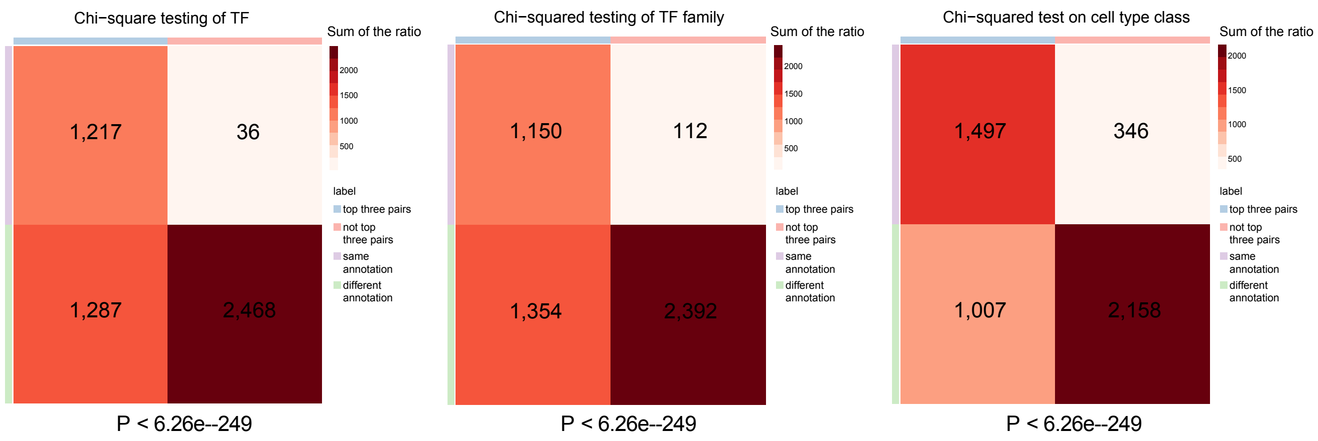


Figure S5. Chi-square testing of TF, TF family, and cell type class. A. Chi-square testing of TF. The color of the heat map indicates the summed ratio in each group. We summed the same or different annotation ratios in the top three or not top three neighboring sample pairs defined by the *k*-sim Pearson, respectively. The color label indicates each group. B and C. Chi-square testing of TF family and cell type class. The heatmap color and the color label indicate the summed ratios and groups similar to A, respectively.

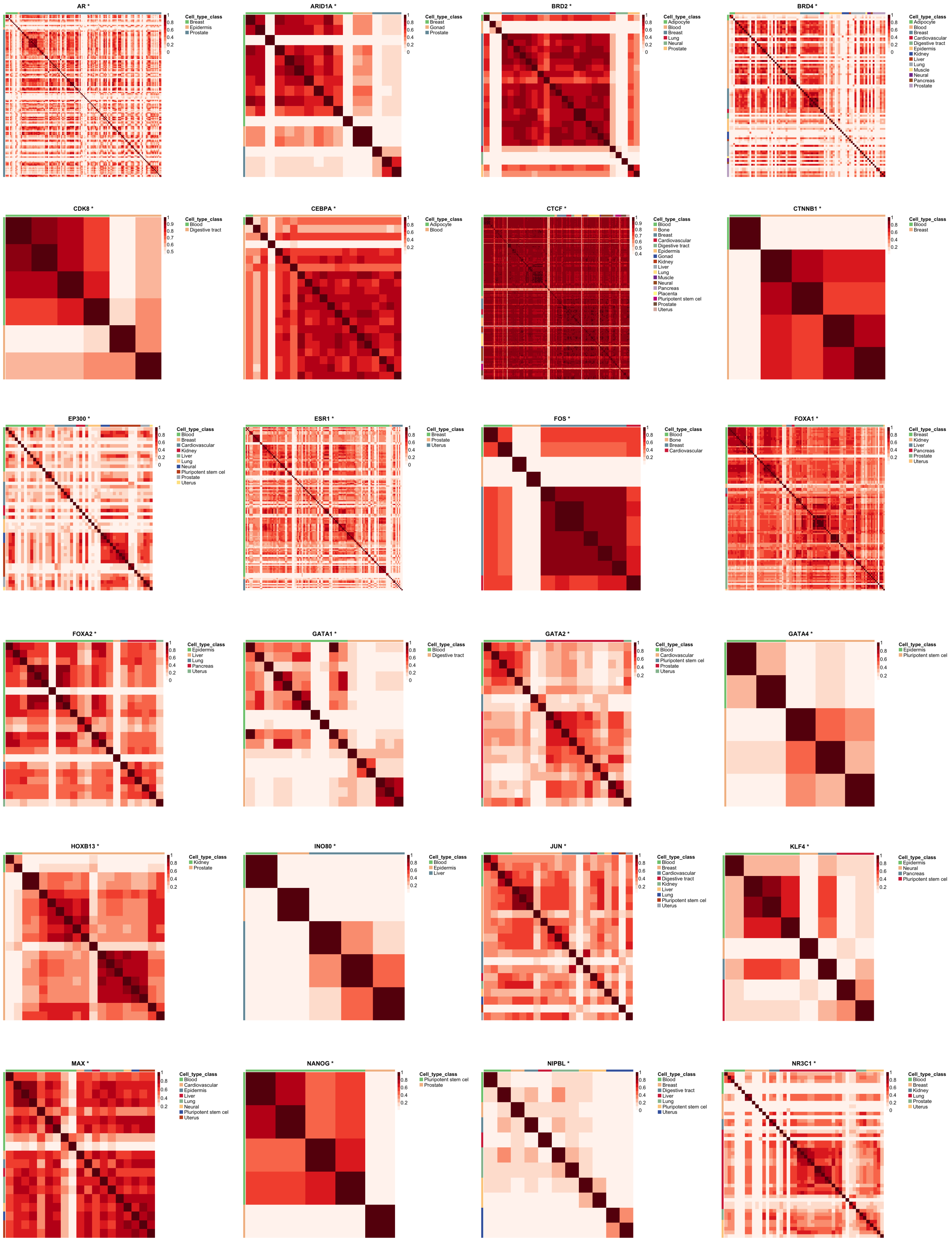

AR *
Cell_type_class
Breast
Epidermis
Prostate
ARID1A *
Cell_type_class
Breast
Gonad
Prostate
BRD2 *
Cell_type_class
Adipocyte
Blood
Breast
Lung
Neural
Prostate
BRD4 *
Cell_type_class
Adipocyte
Blood
Breast
Cardiovascular
Digestive tract
Epidermis
Kidney
Liver
Lung
Muscle
Neural
Pancreas
Prostate
CDK8 *
Cell_type_class
Blood
Digestive tract
CEBPA *
Cell_type_class
Adipocyte
Blood
CTCF *
Cell_type_class
Blood
Bone
Breast
Cardiovascular
Digestive tract
Epidermis
Gonad
Kidney
Liver
Lung
Muscle
Neural
Pancreas
Placenta
Pluripotent stem cel
Prostate
Uterus
CTNNB1 *
Cell_type_class
Blood
Breast
EP300 *
Cell_type_class
Blood
Breast
Cardiovascular
Kidney
Liver
Lung
Neural
Pluripotent stem cel
Prostate
Uterus
ESR1 *
Cell_type_class
Breast
Prostate
Uterus
FOS *
Cell_type_class
Blood
Bone
Breast
Cardiovascular
FOXA1 *
Cell_type_class
Breast
Kidney
Liver
Pancreas
Prostate
Uterus
FOXA2 *
Cell_type_class
Epidermis
Liver
Lung
Pancreas
Uterus
GATA1 *
Cell_type_class
Blood
Digestive tract
GATA2 *
Cell_type_class
Blood
Cardiovascular
Pluripotent stem cel
Prostate
Uterus
GATA4 *
Cell_type_class
Epidermis
Pluripotent stem cel
HOXB13 *
Cell_type_class
Kidney
Prostate
INO80 *
Cell_type_class
Blood
Epidermis
Liver
JUN *
Cell_type_class
Blood
Breast
Cardiovascular
Digestive tract
Kidney
Liver
Lung
Pluripotent stem cel
Uterus
KLF4 *
Cell_type_class
Epidermis
Neural
Pancreas
Pluripotent stem cel
MAX *
Cell_type_class
Blood
Cardiovascular
Epidermis
Liver
Lung
Neural
Pluripotent stem cel
Uterus
NANOG *
Cell_type_class
Pluripotent stem cel
Prostate
NIPBL *
Cell_type_class
Blood
Breast
Digestive tract
Liver
Lung
Pluripotent stem cel
Uterus
NR3C1 *
Cell_type_class
Blood
Breast
Kidney
Lung
Prostate
Uterus

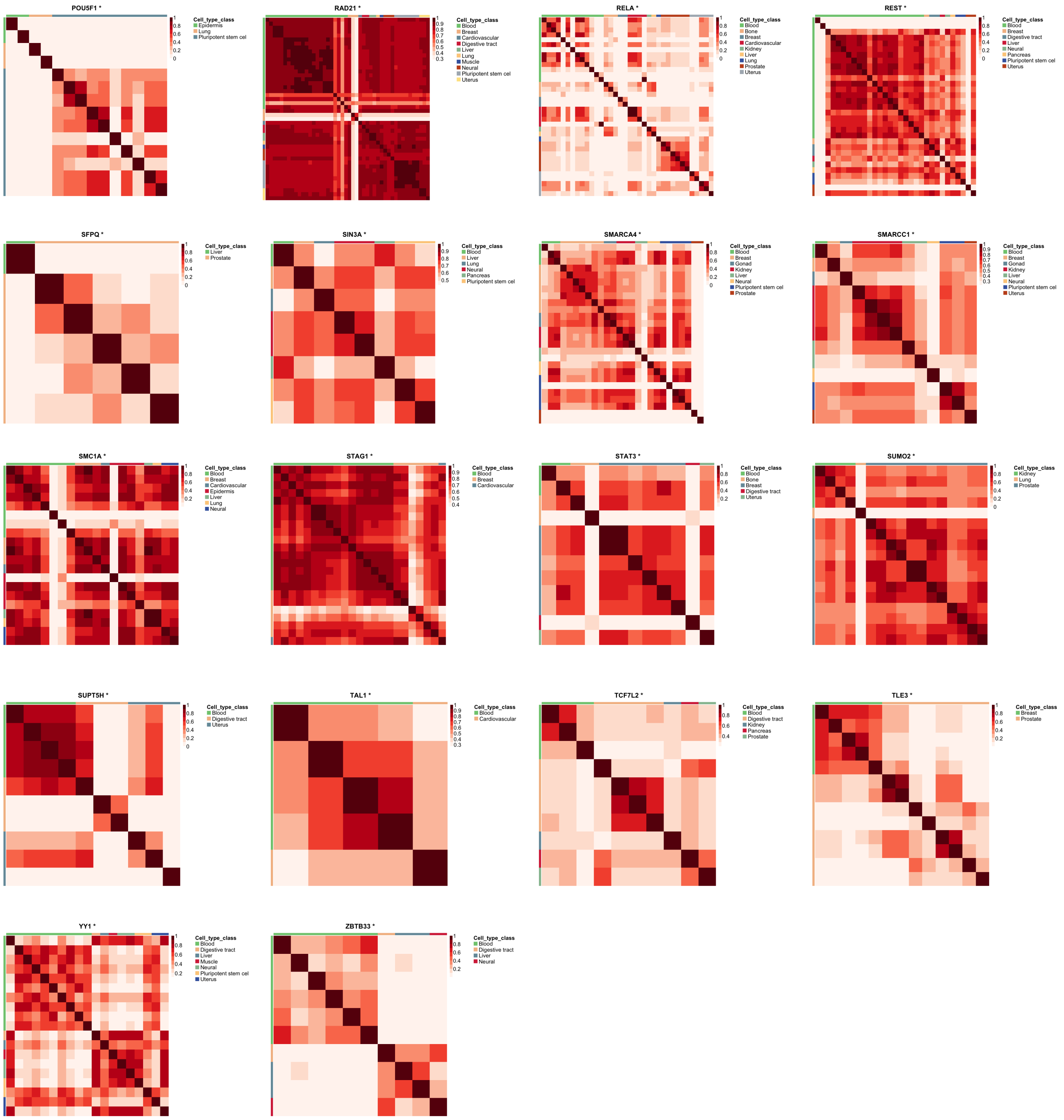


Figure S6. Heat maps of all of the cell type-dependent TFs. The heat map color indicates the *k*-sim Jaccard. The color label of the heat maps indicates the cell type classes. Asterisks indicate statistical significance of ChIP-seq samples with the same and different cell type classes (Mann–Whitney U test, $p < 0.05$).

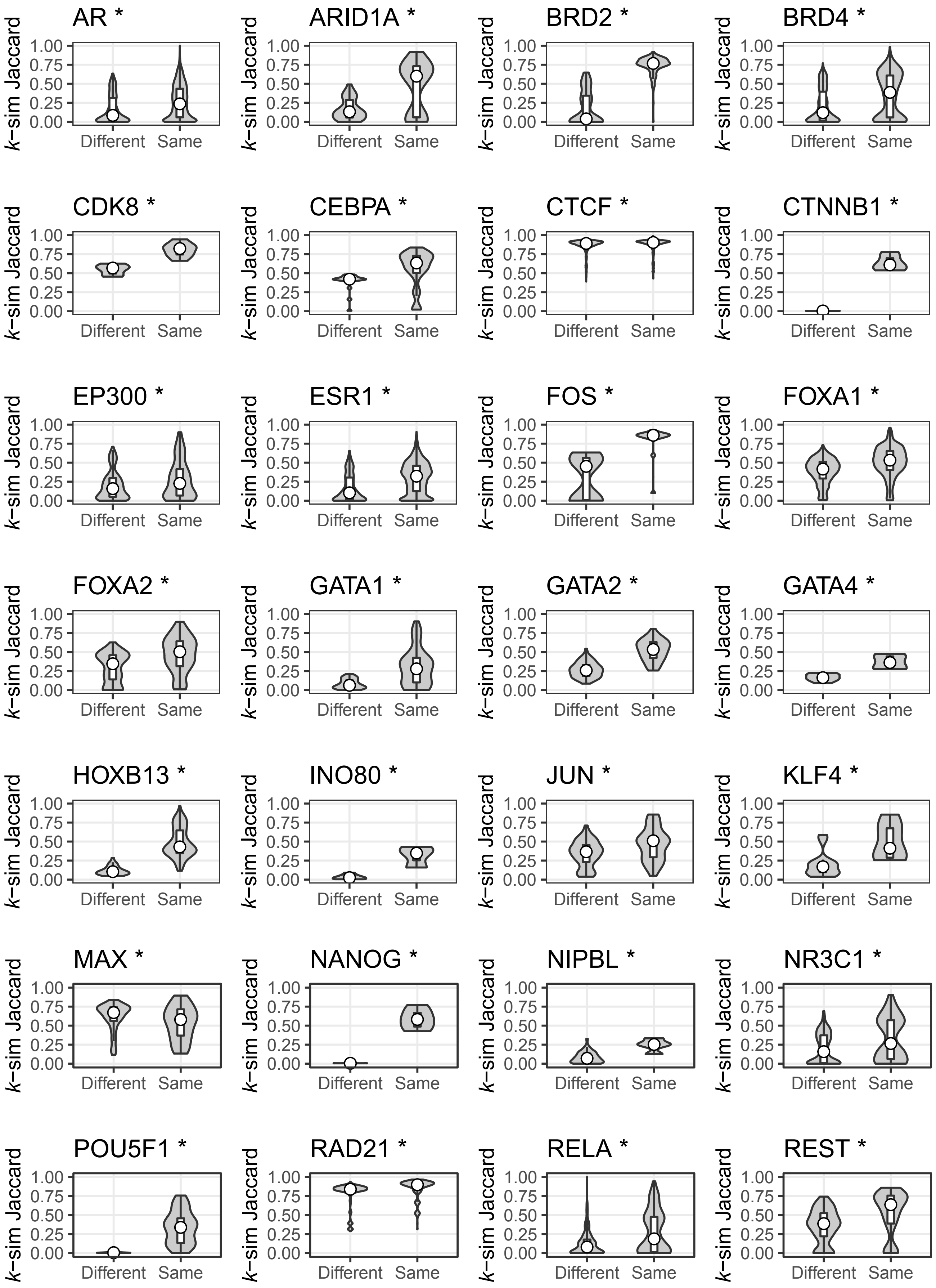

AR *
ARID1A *
BRD2 *
BRD4 *
CDK8 *
CEBPA *
CTCF *
CTNNB1 *
EP300 *
ESR1 *
FOS *
FOXA1 *
FOXA2 *
GATA1 *
GATA2 *
GATA4 *
HOXB13 *
INO80 *
JUN *
KLF4 *
MAX *
NANOG *
NIPBL *
NR3C1 *
POU5F1 *
RAD21 *
RELA *
REST *
k−sim Jaccard
0.00
0.25
0.50
0.75
1.00
Different
Same

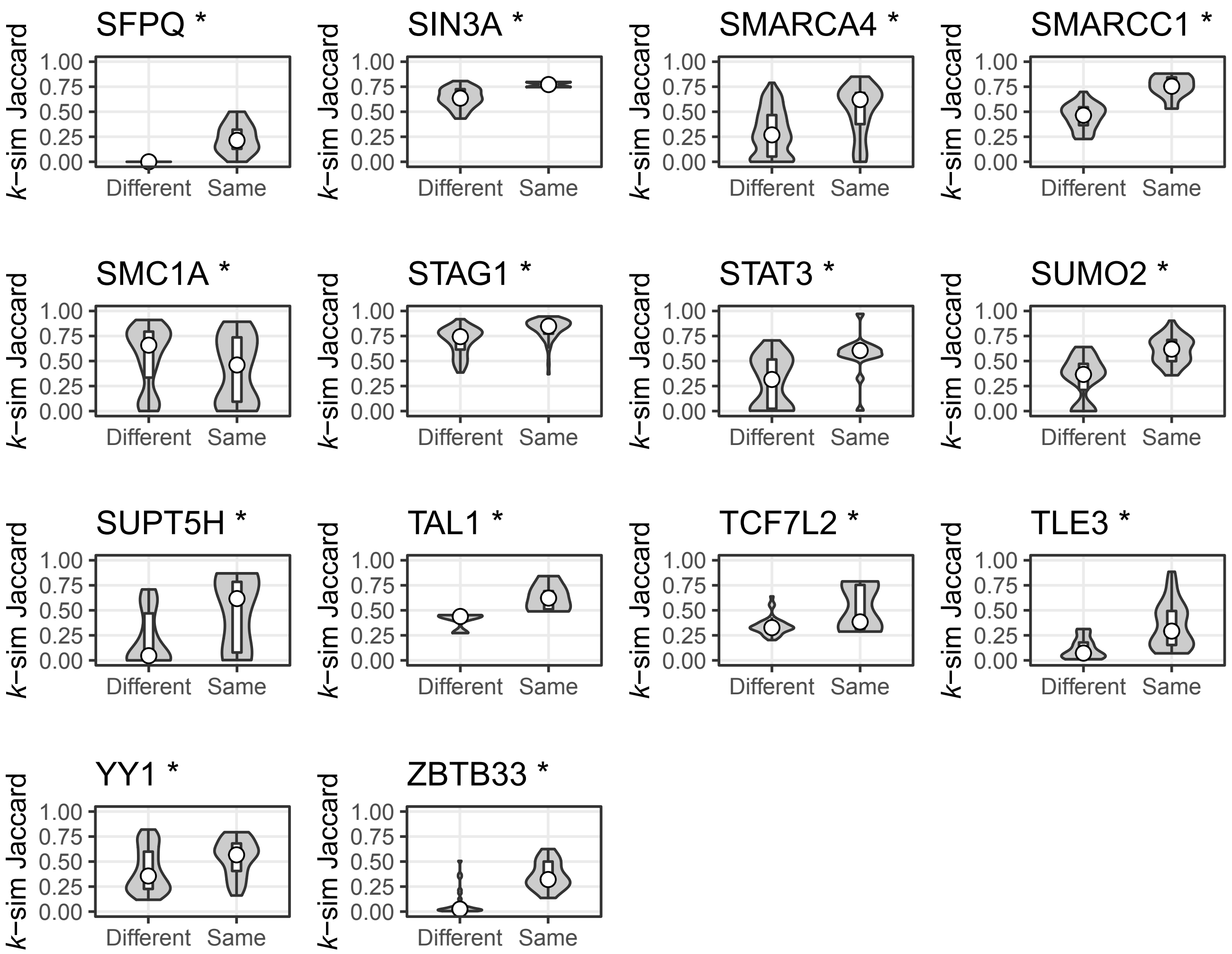


Figure S7. Violin plots of all cell type-dependent TFs. The y-axis indicates the*k*-sim Jaccard. The same and different groups are arranged along the x-axis. Asterisks indicate the statistical significance of the ChIP-seq samples with the same and different cell type classes (Mann–Whitney U test, $p < 0.05$)

Tahara *et al*., Figure S7

**A**

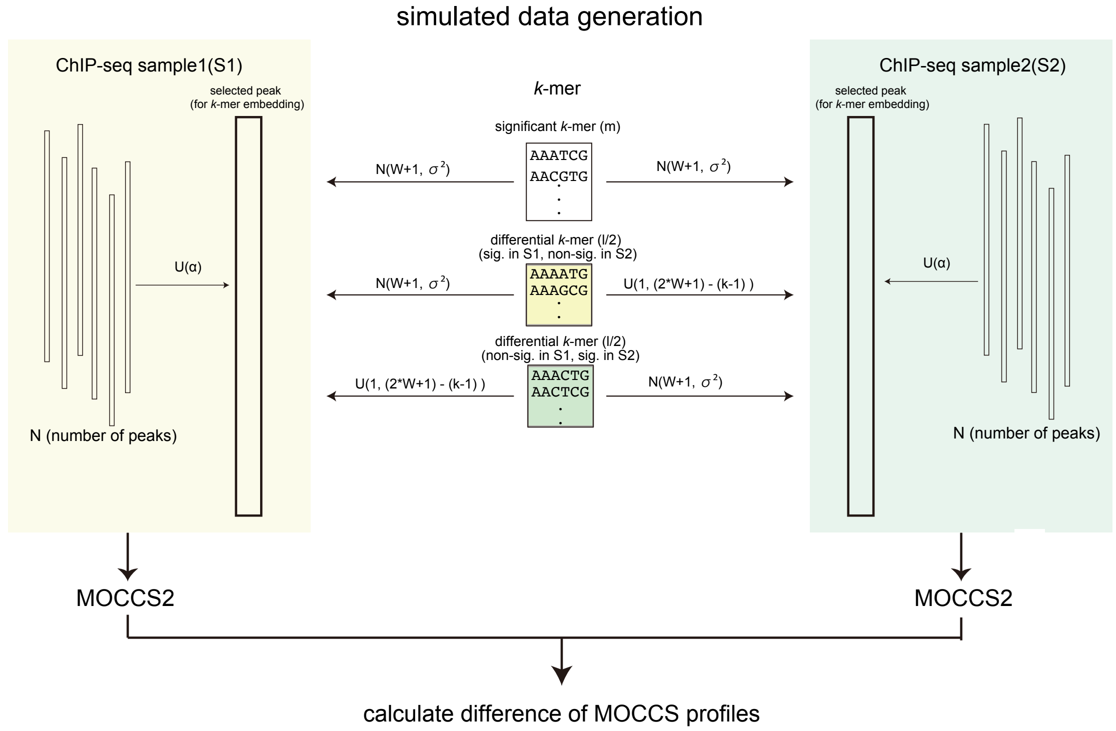


**B**

| simulation | α | N | σ |
|---|---|---|---|
| #1 | [0.10, 0.20] | 12000 | W/5 |
| #2 | [0.01, 0.05] | 12000 | W/5 |
| #3 | [0.05, 0.10] | 12000 | W/5 |
| #4 | [0.10, 0.20] | 6000 | W/5 |
| #5 | [0.10, 0.20] | 12000 | W/2 |

**α** : The percentage of the input sequences containing embedded "true significant *k*-mers"

**N** : The number of peaks in a ChIP-seq sample

**σ** : The standard deviation of the embedded "true significant *k*-mers" from the center of the peak

**W**: 2W+1bp means a length of a peak in a ChIP-seq

**l** : Number of embedded differential *k*-mers

**m** : Number of embedded significant *k*-mers

Figure S8. Simulation of differential *k*-mer detection. A. Simulated data processing. We prepared simulated data with an embedded "true differential *k*-mer" and "true significant *k*-mer" by embedding a "true" *k*-mer within α% of a randomly generated sample of $2W + 1 - \text{bp}$ ($W = 350$) DNA sequences and applying MOCCS2. "True significant *k*-mers" were embedded following a normal distribution whose mean was $W + 1$ and whose standard deviation was σ.
"True differential *k*-mers" were embedded in S1 (or S2), the same as "true significant *k*-mers" , and were embedded in S2 (or S1) following a uniform distribution whose mean was 1 and whose standard deviation was $(2 \times W + 1) - (k - 1)$.
B. Parameters in each simulation condition from #1 to #5. L is the number of differential *k*-mers and m is the number of significant *k*-mers.

**A**

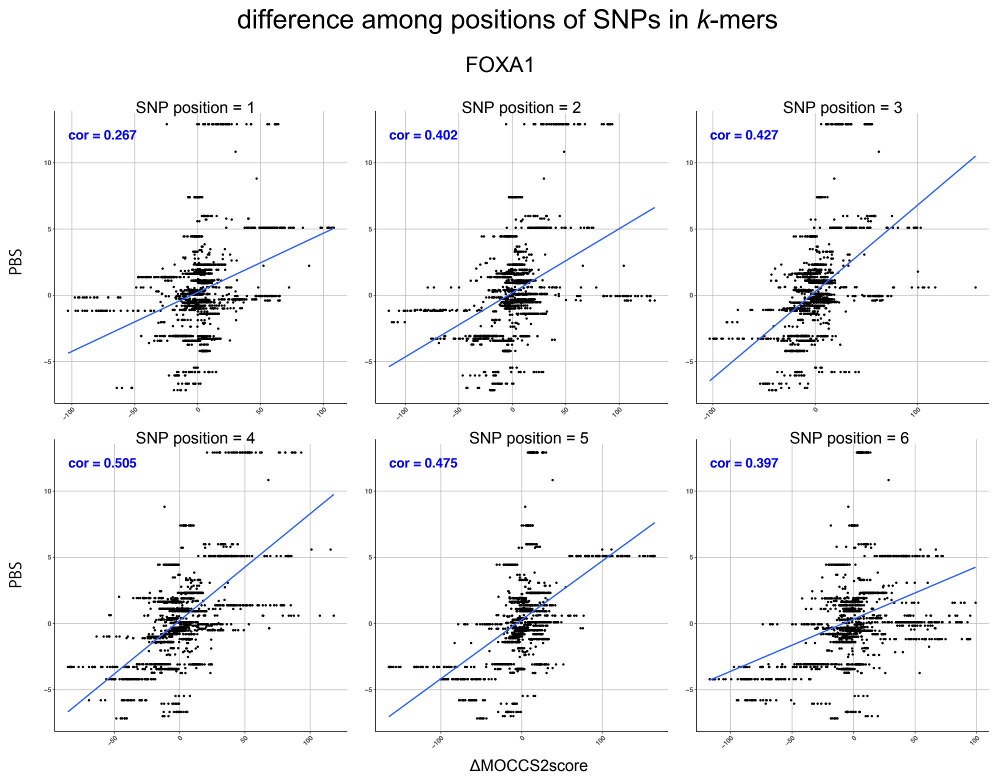


**B**

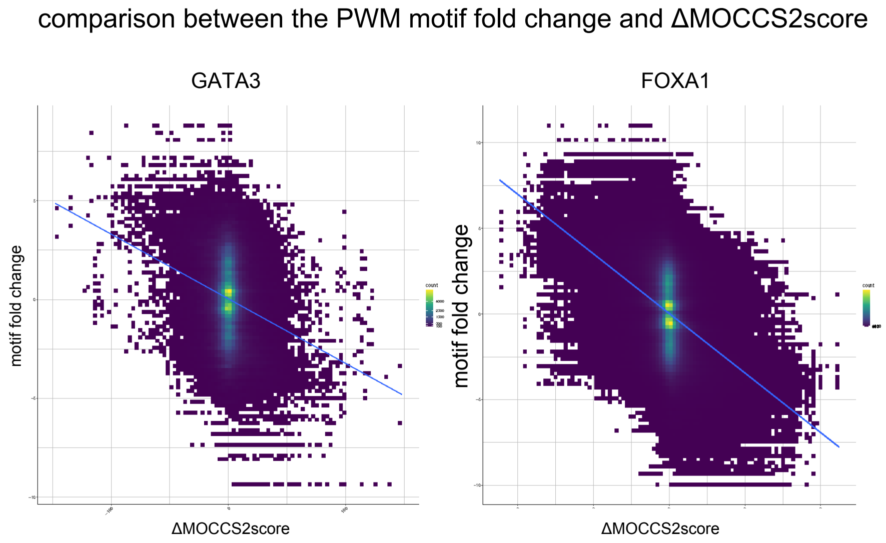


Figure S9. ΔMOCCS2score profiles agree with the *in vitro* SNP-SELEX and PWM motif fold change.

A. Difference in the ΔMOCCS2score profile consistency among the positions of SNPs in *k*-mers. The *k*th SNP position means the *k*th allele from the left in the *k*-mer. B. The ΔMOCCS2score agrees with the PWM motif fold change.

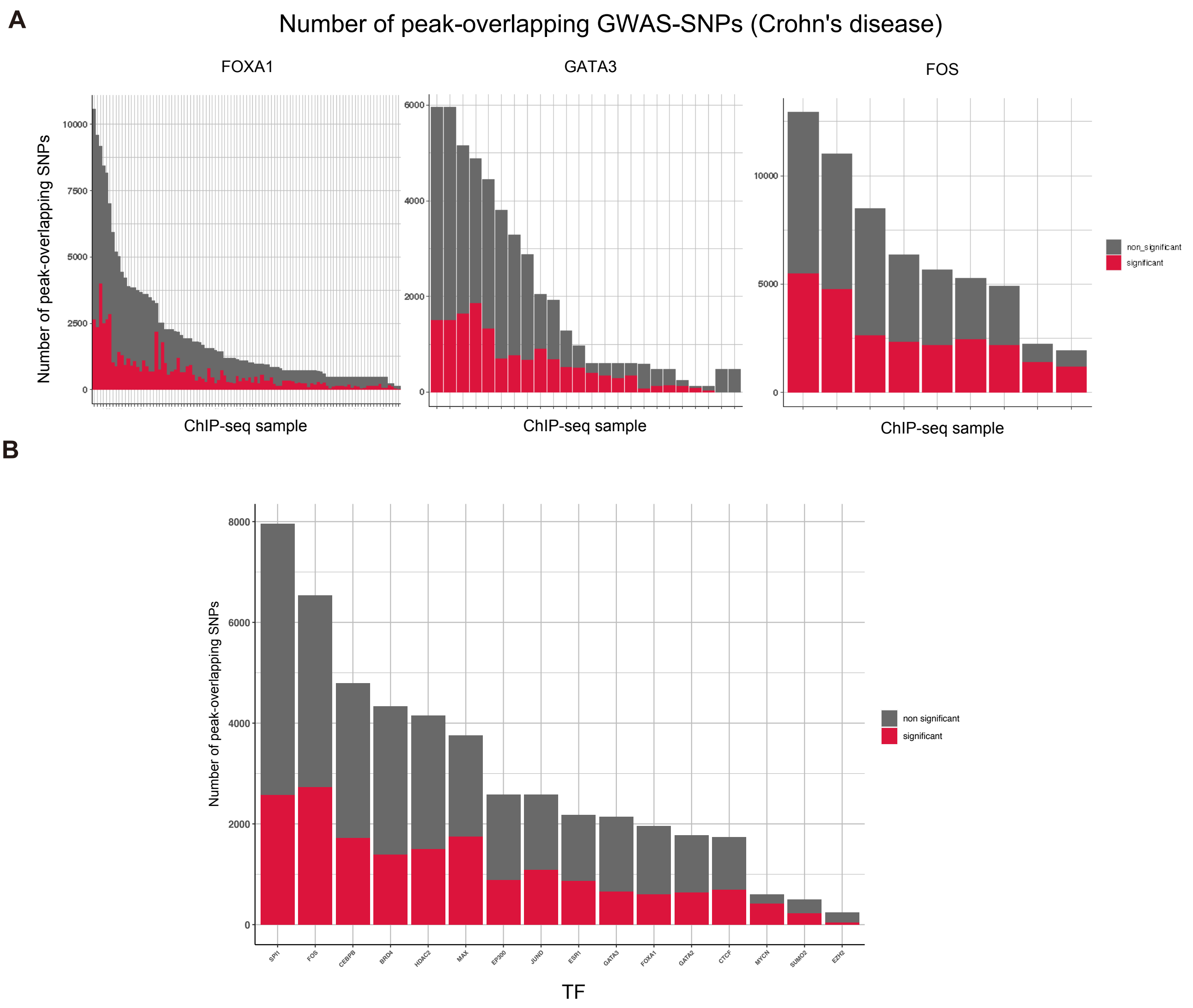


Figure S10. Number of differential peak-overlapping GWAS-SNPs. A. Number of peak-overlapping GWAS-SNPs with significant ΔMOCCS2scores of Crohn's disease in one ChIP-seq sample. The red fraction shows the number of GWAS-SNPs with significant ΔMOCCS2scores ($q < 0.05$). B. Number of differential peak-overlapping GWAS-SNPs of Crohn's disease in the ChIP-seq samples of the same TFs.

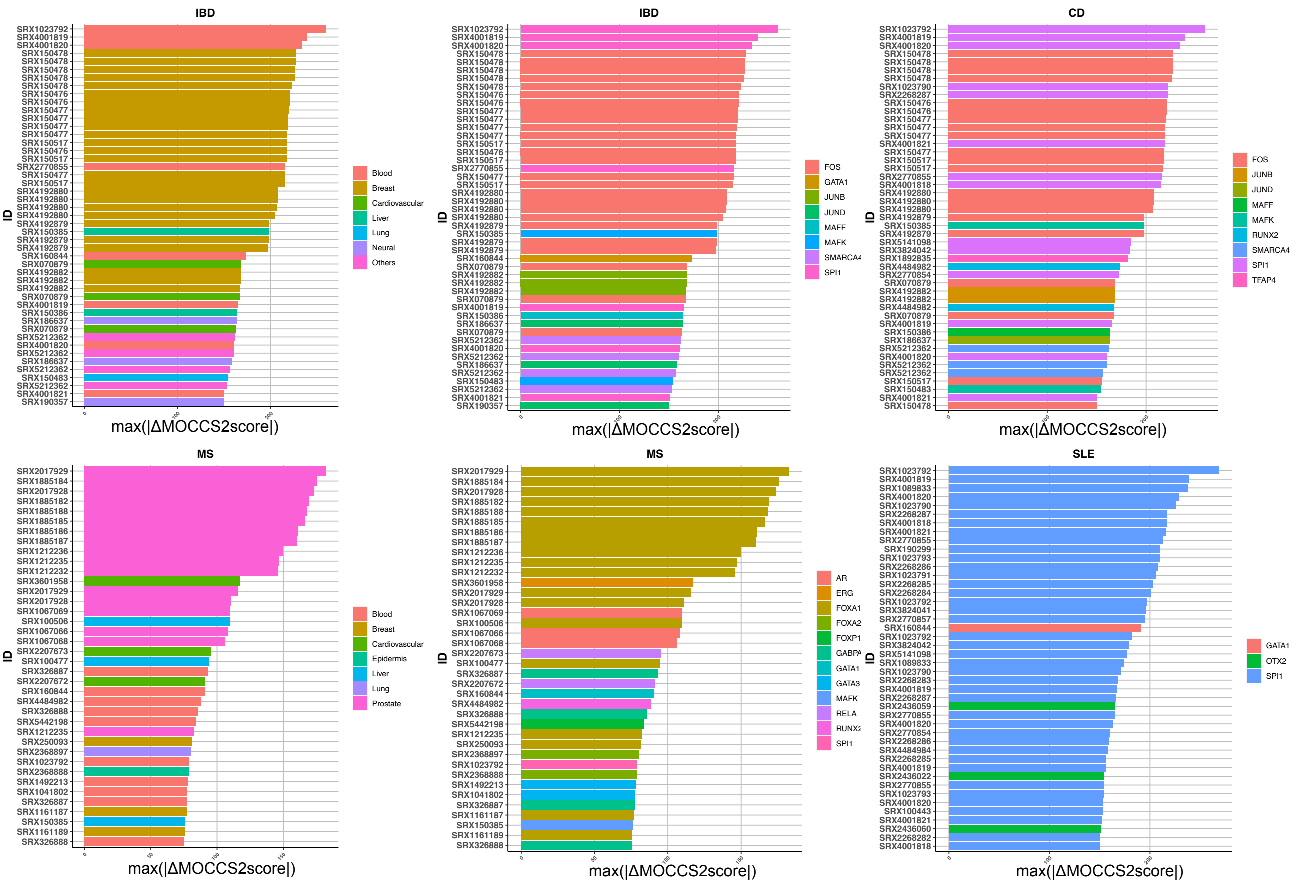


Figure S11. Prediction of SNP-affected TFs and cell type classes through ΔMOCCS2score profiles. Top ChIP-seq samples with high ΔMOCCS2scores in each phenotype (IBD, inflammatory bowel disease; CD, Crohn's disease; MS, multiple sclerosis; SLE, systemic lupus erythematosus). Each SNP in a ChIP-seq sample has a ΔMOCCS2score, and the SNPs have been arranged by the ΔMOCCS2score. Each bar is each SNP from a ChIP-seq sample. Bar graph colors show TFs or cell type class.

Tahara *et al.*, Figure S11

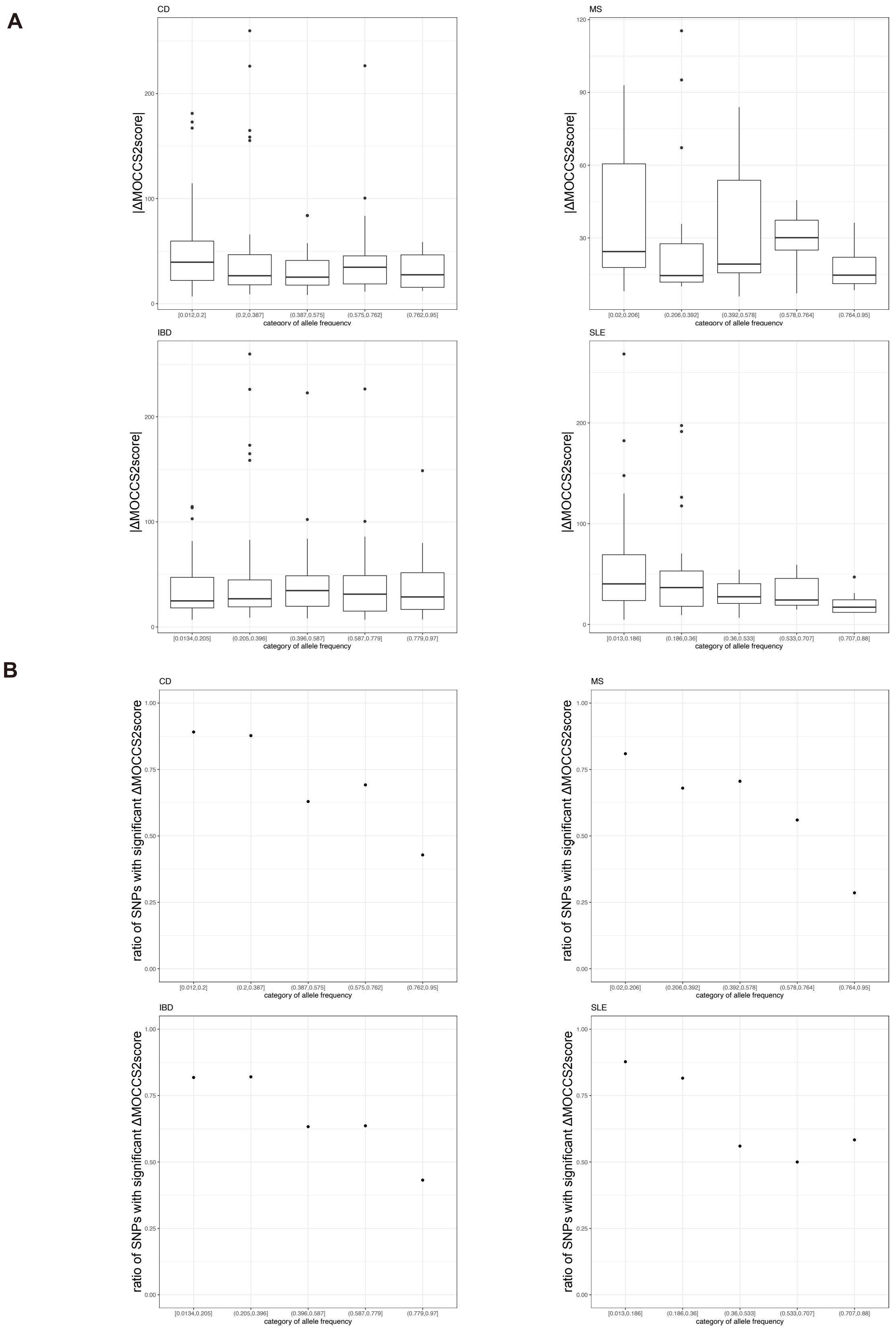


Figure. S12. Association between the allele frequency and ΔMOCCS2score. A. Association between the allele frequency and the absolute values of the ΔMOCCS2score. B. Association between the allele frequency and the ratio of SNPs with a significant ΔMOCCS2score.

Tahara *et al.*, Figure S12
